# They call me the wanderer – Neurovascular anatomy of dwarfed dinosaur implies precociality in sauropods

**DOI:** 10.1101/2022.08.01.502289

**Authors:** Marco Schade, Nils Knötschke, Marie K. Hörnig, Carina Paetzel, Sebastian Stumpf

## Abstract

Macronaria, a group of mostly colossal sauropod dinosaurs, comprised the largest terrestrial vertebrates of Earth’s history. However, some of the smallest sauropods belong to this group as well. The Late Jurassic macronarian *Europasaurus holgeri* is one of the best-known sauropods worldwide. So far, the braincase material of this taxon from Germany pended greater attention. With the aid of microCT, we report on the neuroanatomy of the almost complete braincase of an adult individual, as well as the inner ears of one other adult and several juveniles (also containing so far unknown vascular cavities). The presence of large and morphologically adult inner ears in juvenile material suggests precociality. Our findings add to the diversity of neurovascular anatomy in sauropod braincases and buttress the perception of sauropods as fast-growing and autonomous giants with manifold facets of reproductive and social behavior. This suggests that – apart from sheer size – little separated the island dwarf *Europasaurus* from its large-bodied relatives.

## Introduction

Sauropoda is a taxon of saurischian dinosaurs and comprise popular taxa like *Diplodocus*, *Giraffatitan* and *Argentinosaurus*^1^. Sauropods were diverse and successfully distributed worldwide (e.g.^1, 2^).The oldest indubitable sauropod taxa are known from Early Jurassic strata; the youngest representatives went extinct during the end-Cretaceous disaster^2^. Whereas bipedal early sauropodomorphs (in which sauropods are phylogenetically nested) were probably capable of swiftly tracking down prey^3^, the later evolutionary history of the group is characterized by an unrivaled increase in body size (among land-dwelling vertebrates), accompanied with herbivory, an extreme elongation in neck length and graviportal quadrupedality (e.g.^1, 4, 5^).

While fossil braincases are generally rare, studies of sauropod endocrania are surprisingly numerous (e.g. ^6,7,8^), serving as a good base for comparisons. Potentially, aspects of lifestyle can be inferred from certain morphological details of cavities that once contained the brain, inner ear and other associated neurovascular structures within the bony braincase of vertebrates (e.g.^9,10,11,12^). Furthermore, ontogenetically induced morphological shifts of neuroanatomy can hint towards different ecological tendencies within a species, for example, in respect to bipedal or quadrupedal locomotion^13^.

The middle Kimmeridgian (Late Jurassic) sauropod *Europasaurus* (represented by a single species, *E*. *holgeri*) is regarded as an unequivocal example for insular dwarfism with paedomorphic features, having reached adult body lengths of nearly 6 metres and weighing about 800 kg^14,15,16,17^. From this taxon, a great number of cranial and postcranial fossil bones are known (housed in the Dinosaurier-Freilichtmuseum Münchehagen/Verein zur Förderung der Niedersächsischen Paläontologie e.V., Rehburg - Loccum, Münchehagen, Germany), of which the former hint to at least 14 individuals of different ontogenetic stages^17^.

The paratype specimen of *Europasaurus*, DFMMh/FV 581.1, comprises a largely complete, articulated and probably mature braincase, with DFMMh/FV 581.2 & 3 representing the respective detached parietals (Figs. 1–3; Supplementary Figs. 1–4). The outer morphology of this material has previously been described^17^. For this study, the parietals were put in their place on the remaining endocranium and subsequently documented with microCT. The endocranial cavities which once housed the brain, inner ears and other soft neuroanatomical structures, such as nerves and blood supply, were manually segmented. The articulated specimens DFMMh/FV 581.1, 2 & 3 measure about 120 mm in mediolateral width, 80 mm anteroposteriorly and 100 mm dorsoventrally.

**Fig. 1:**
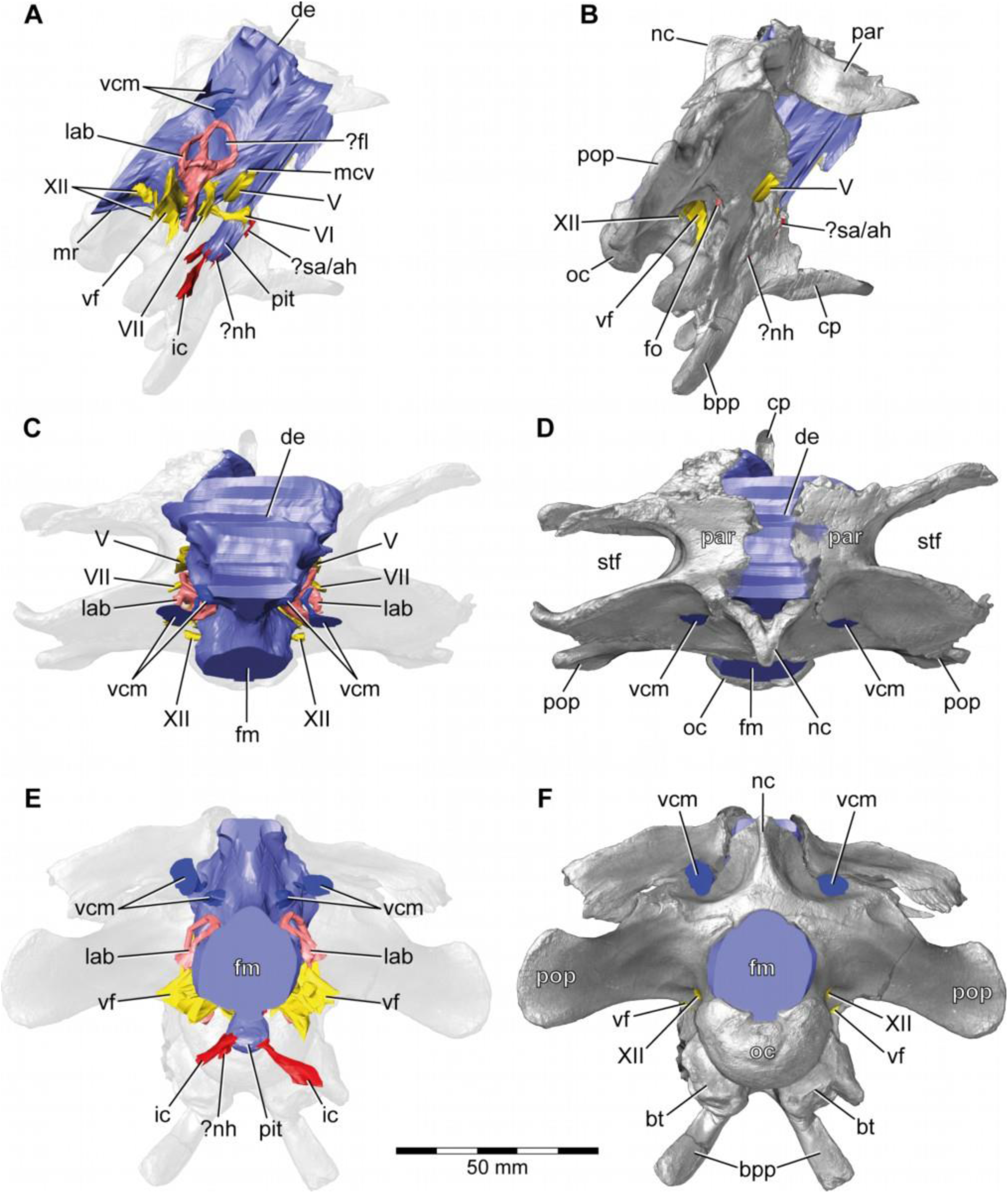
*Europasaurus holgeri*, 3D model of the braincase endocast with endosseous labyrinths and neurovascular canals of DFMMh/FV 581.1, 2 & 3 with transparent (**A**,**C**,**E**) and covering (**B**,**D**,**F**) volume rendering of the bony braincase in (**A**,**B**) right lateral, (**C**,**D**) dorsal and (**E**,**F**) posterior view. Note that scale mainly applies to posterior perspective (**E**,**F**). ?fl, potential floccular recess; ?nh, potential canal for the neurohypophysis; ?sa/ah, potential sphenoidal artery/canal for the adenohypophysis; bpp, basipterygoid process; bt, basal tuber; cp, cultriform process; de, dorsal expansion; ic, internal carotid; fm, foramen magnum; fo, fenestra ovalis; lab, endosseous labyrinth; mcv, mid cerebral vein; mr, medial ridge; nc, sagittal nuchal crest; oc, occipital condyle; par, parietal; pit, pituitary; pop, paroccipital process; stf, supratemporal fenestra; vcm, vena capitis media; vf, vagal foramen; V, trigeminal nerve; VI, abducens nerve; VII, facial nerve; XII, hypoglossal nerve.

**Fig. 2:**
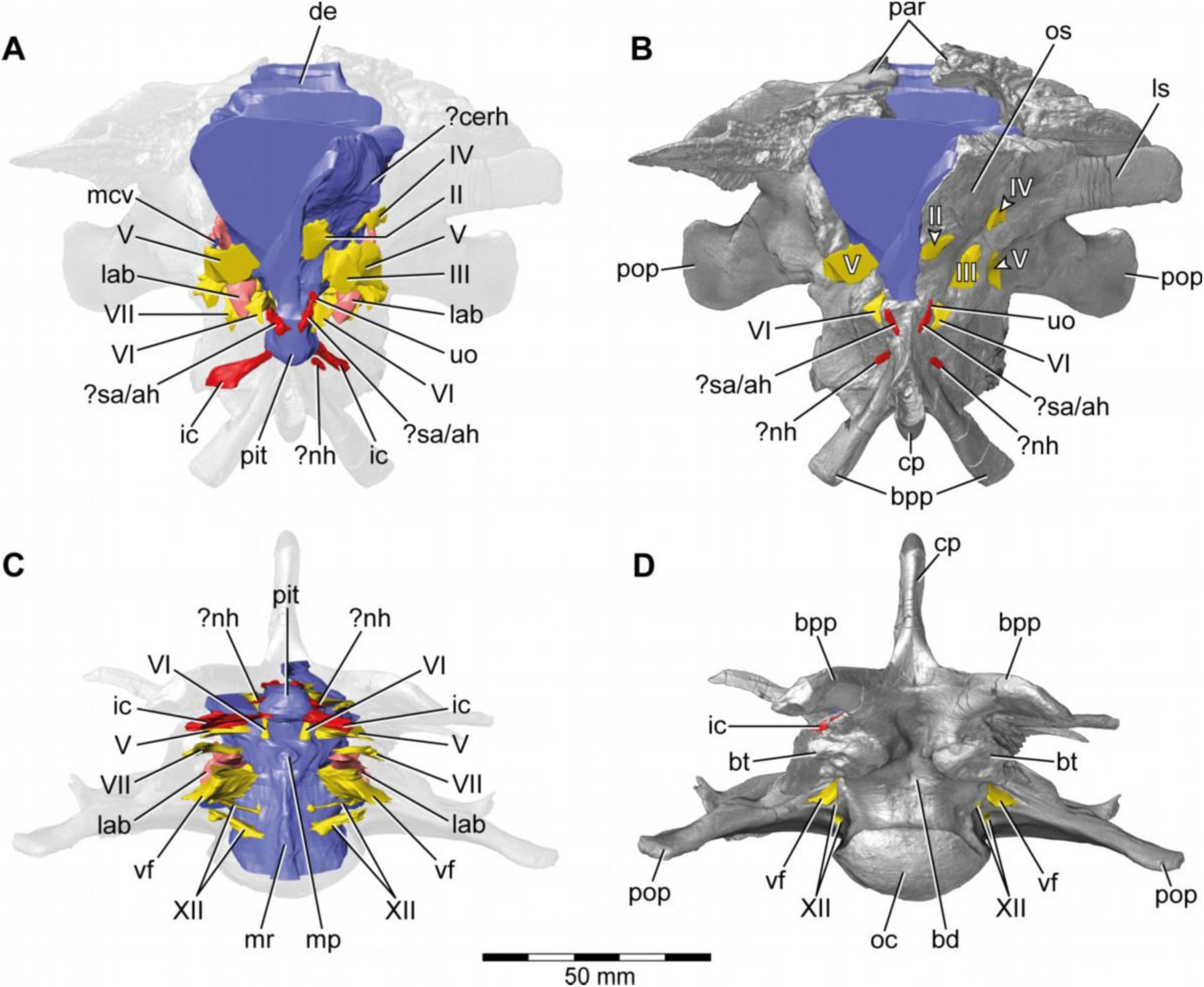
*Europasaurus holgeri*, 3D model of the braincase endocast with endosseous labyrinths and neurovascular canals of DFMMh/FV 581.1, 2 & 3 with transparent (**A**,**C**) and covering (**B**,**D**) volume rendering of the bony braincase in (**A**,**B**) anterior (**C**,**D**) and ventral view. Note that scale mainly applies to ventral perspective (**C**,**D**). ?cerh, potential cerebral hemisphere; ?nh, potential canal for the neurohypophysis; ?sa/ah, potential sphenoidal artery/canal for the adenohypophysis; bd, blind depression; bpp, basipterygoid process; bt, basal tuber; cp, cultriform process; de, dorsal expansion; ic, internal carotid; fm, foramen magnum; lab, endosseous labyrinth; ls; laterosphenoid; mcv, mid cerebral vein; mp, median protuberance; mr, medial ridge; oc, occipital condyle; os, orbitosphenoid; par, parietal; pit, pituitary; pop, paroccipital process; uo, unclear opening; vf, vagal foramen; II, optic nerve; III, oculomotor nerve; IV, trochlear nerve; V, trigeminal nerve; VI, abducens nerve; VII, facial nerve; XII, hypoglossal nerve.

**Fig. 3:**
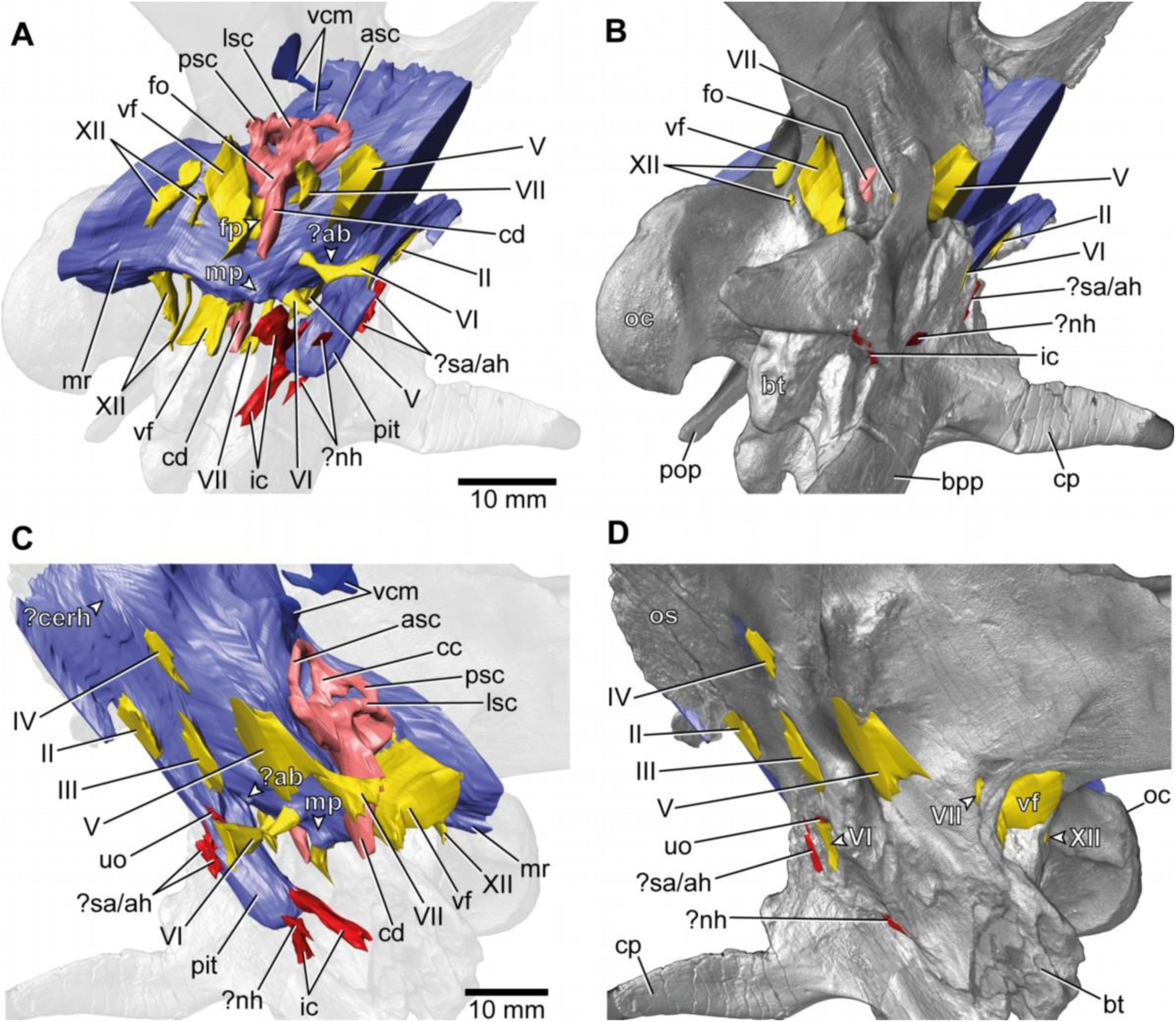
*Europasaurus holgeri*, 3D model of the braincase endocast with endosseous labyrinths and neurovascular canals of DFMMh/FV 581.1, 2 & 3 with transparent (**A**,**C**) and covering (**B**,**D**) volume rendering of the bony braincase in (**A**,**B**) right ventrolateral and (**C**,**D**) left lateral view. ?ab, potential basilar artery; ?cerh, potential cerebral hemisphere; ?nh, potential canal for the neurohypophysis; ?sa/ah, potential sphenoidal artery/canal for the adenohypophysis; bpp, basipterygoid process; bt, basal tuber; cp, cultriform process; ic, internal carotid; fo, fenestra ovalis; fp, fenestra pseudorotunda; mp, median protuberance; mr, medial ridge; oc, occipital condyle; os, orbitosphenoid; pit, pituitary; pop, paroccipital process; uo, unclear opening; vcm, vena capitis media; vf, vagal foramen; II, optic nerve; III, oculomotor nerve; IV, trochlear nerve; V, trigeminal nerve; VI, abducens nerve; VII, facial nerve; XII, hypoglossal nerve.

**Fig. 4:**
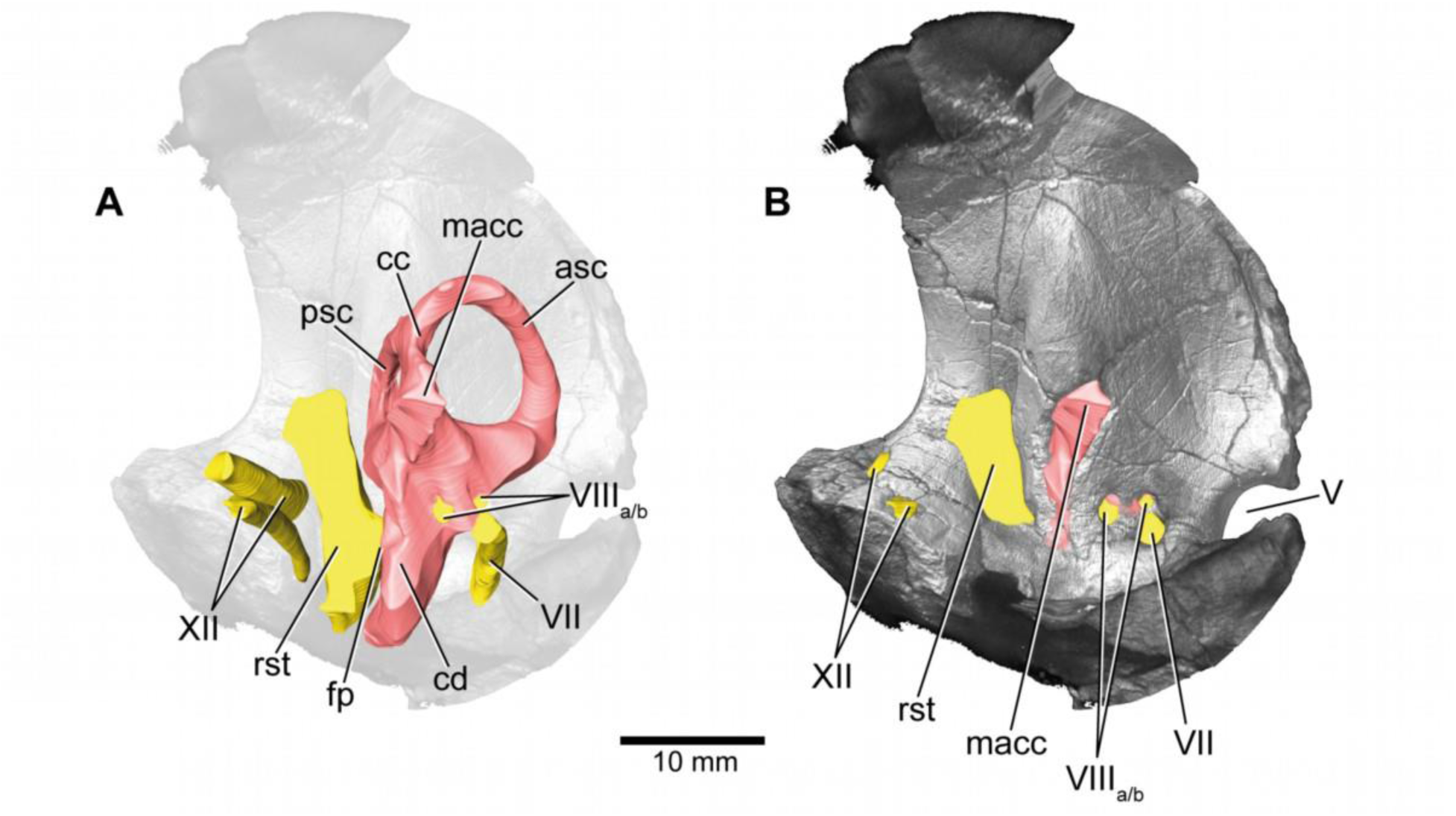
*Europasaurus holgeri*, 3D model of the left endosseous labyrinth region in DFMMh/FV 1077 with transparent (**A**) and covering (**B**) volume rendering of the bony braincase in medial view. asc, anterior semicircular canal; cc, common crus; cd, cochlear duct; fp, fenestra pseudorotunda; lsc, lateral semicircular canal; macc, medial aspect of common crus; psc, posterior semicircular canal; rst, recessus scalae tympani; V, trigeminal nerve opening; VII, facial nerve; VIIIa/b, both branches of the vestibulocochlear nerve; XII, hypoglossal nerve.

Additionally, the specimens DFMMh/FV 1077 (Fig. 4; Supplementary Figs. 5, 6; adult fragmentary braincase, complete inner ear), DFMMh/FV 466+205 (Figs. 5, 6; Supplementary Figs. 7–11; juvenile prootic and otoccipital, nearly complete inner ear; the common bond of these two specimens has not been recognized in former studies^17^), DFMMh/FV 964 and DFMMh/FV 561 (Fig. 7; Supplementary Figs. 8, 9; prootics of uncertain maturity, anterior labyrinth), DFMMh/FV 981.2, DFMMh/FV 898 and DFMMh/FV 249 (Fig. 8; Supplementary Figs. 10, 11; juvenile otoccipitals, posterior labyrinth) were documented with microCT. Since the isolated specimens contain different parts of the endosseous labyrinths, cranial nerves and vascular cavities, the respective digital models were reconstructed in order to describe, compare and contextualize their characteristics. Whereas the smallest of these specimens (DFMMh/FV 898) hints to an approximate posterior skull width of under 5 cm, the largest specimens DFMMh/FV 581.1 and DFMMh/FV 1077 suggest a maximum mediolateral width of about 14 cm.

**Fig. 5:**
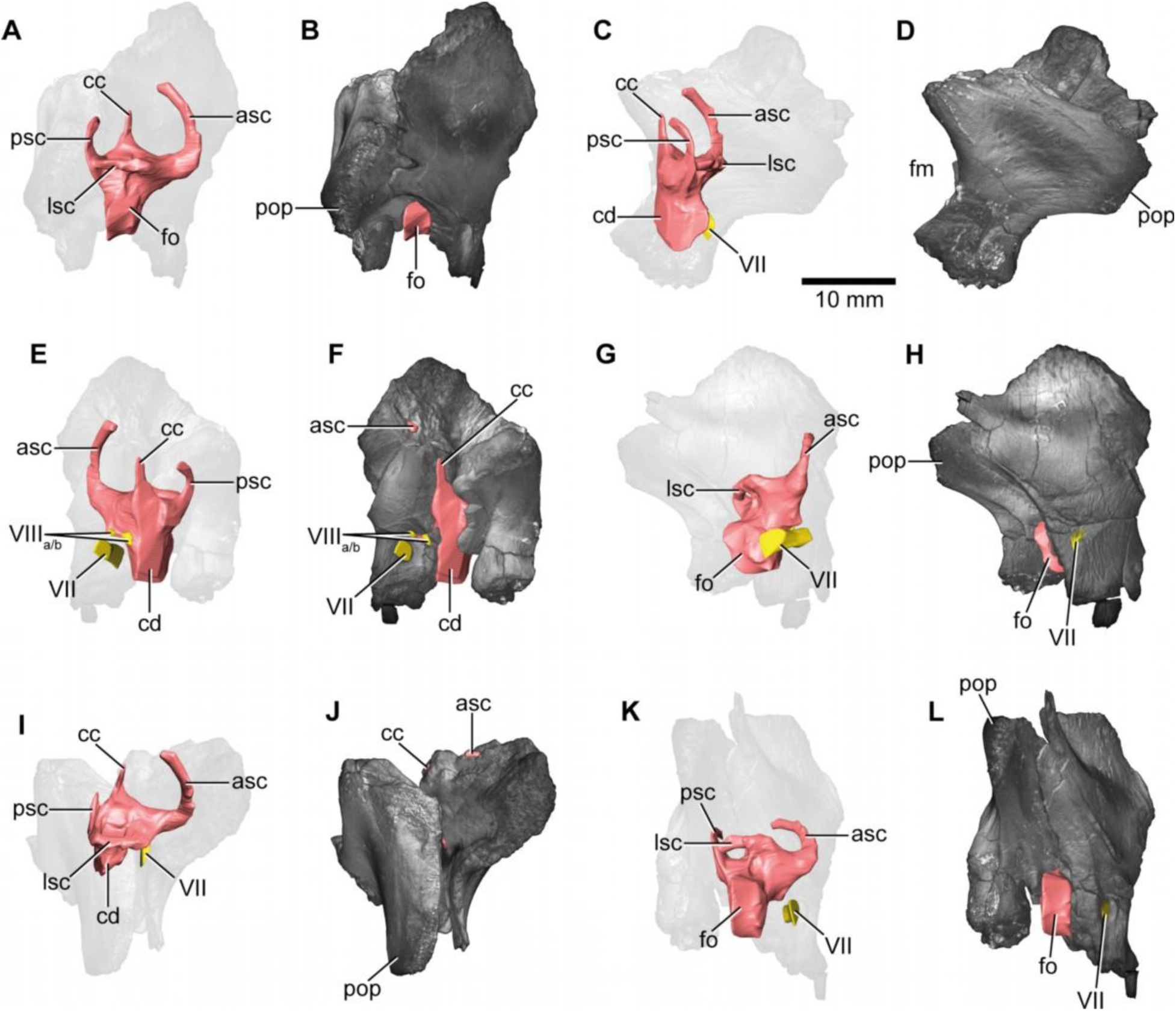
*Europasaurus holgeri*, 3D model of the right endosseous labyrinth in DFMMh/FV 466+205 with transparent (**A**,**C**,**E**,**G**,**I**,**K**) and covering (**B**,**D**,**F**,**H**,**J**,**L**) volume rendering of the bony braincase remains in (**A**,**B**) lateral, (**C**,**D**) posterior, (**E**,**F**) medial, (**G**,**H**) anterolateroventral, (**I**,**J**) dorsolateral and (**K**,**L**) lateroventral view; in respect to the endosseous labyrinth. Note that scale mainly applies to posterior perspective (**C**,**D**), and that VII and VIIIa/b are not shown in **A** and **B**. asc, anterior semicircular canal; cc, common crus; cd, cochlear duct; fm, foramen magnum; fo, fenestra ovalis; lsc, lateral semicircular canal; pop, paroccipital process; psc, posterior semicircular canal; VII, facial nerve; VIIIa/b, both branches of the vestibulocochlear nerve.

**Fig. 6:**
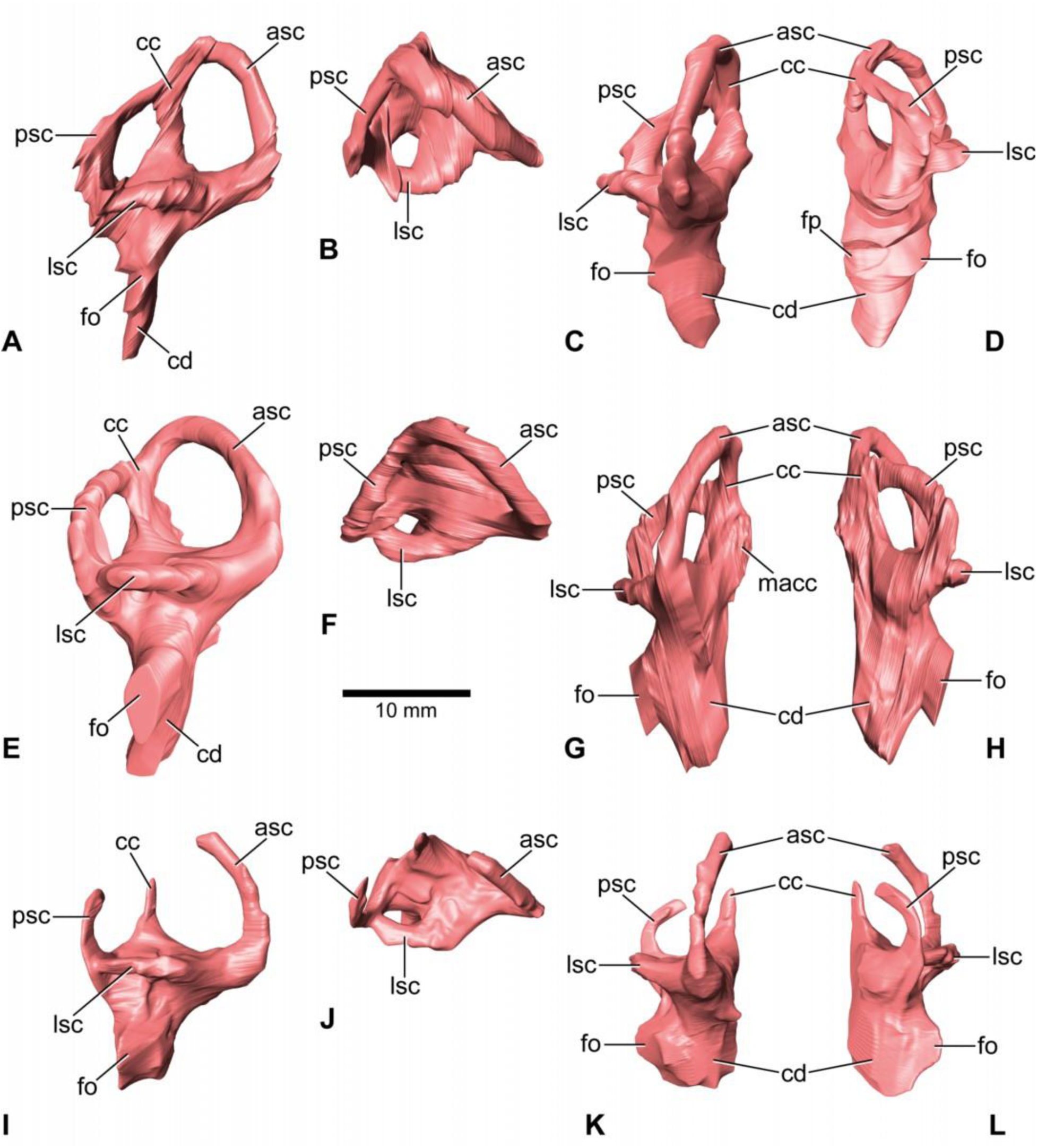
*Europasaurus holgeri*, 3D models of the endosseous labyrinth of DFMMh/FV 581.1 (**A**- **D**), DFMMh/FV 1077 (**E**-**H**; note that this model is mirrored) and DFMMh/FV 466+205 (**I**-**L**) in (**A**,**E**,**I**) lateral, (**B**,**F**,**J**) dorsal, (**C**,**G**,**K**) anterior and (**D**,**H**,**L**) posterior view. Note that scale mainly applies to dorsal perspective (**B**,**F**,**J**). asc, anterior semicircular canal; cc, common crus; cd, cochlear duct; fo, fenestra ovalis; fp, fenestra pseudorotunda; lsc, lateral semicircular canal; macc, medial aspect of common crus; psc, posterior semicircular canal.

**Fig. 7:**
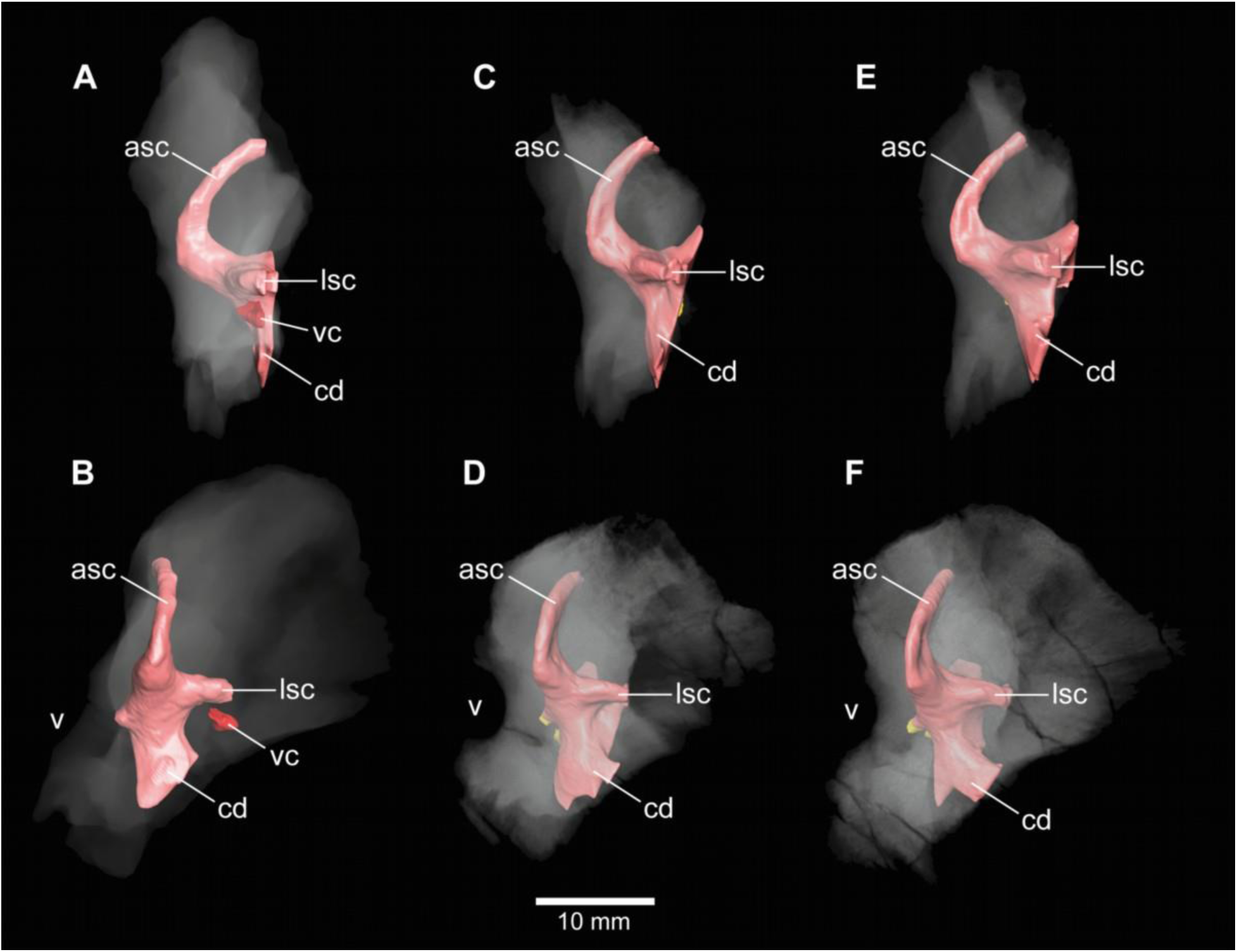
*Europasaurus holgeri*, 3D models of the anterior portions of the endosseous labyrinth in (**A**,**B**; note that this model is mirrored) DFMMh/FV 466, (**C**,**D**) DFMMh/FV 561 and (**E**,**F**) DFMMh/FV 964 in (**A**,**C**,**E**) lateral and (**B**,**D**,**F**) anterolateral view; in respect to the endosseous labyrinth. Note that scale mainly applies to anterolateral perspective (**B**,**D**,**F**). asc, anterior semicircular canal; cd, cochlear duct; lsc, lateral semicircular canal; vc, vascular cavity; V, trigeminal nerve opening.

**Fig. 8:**
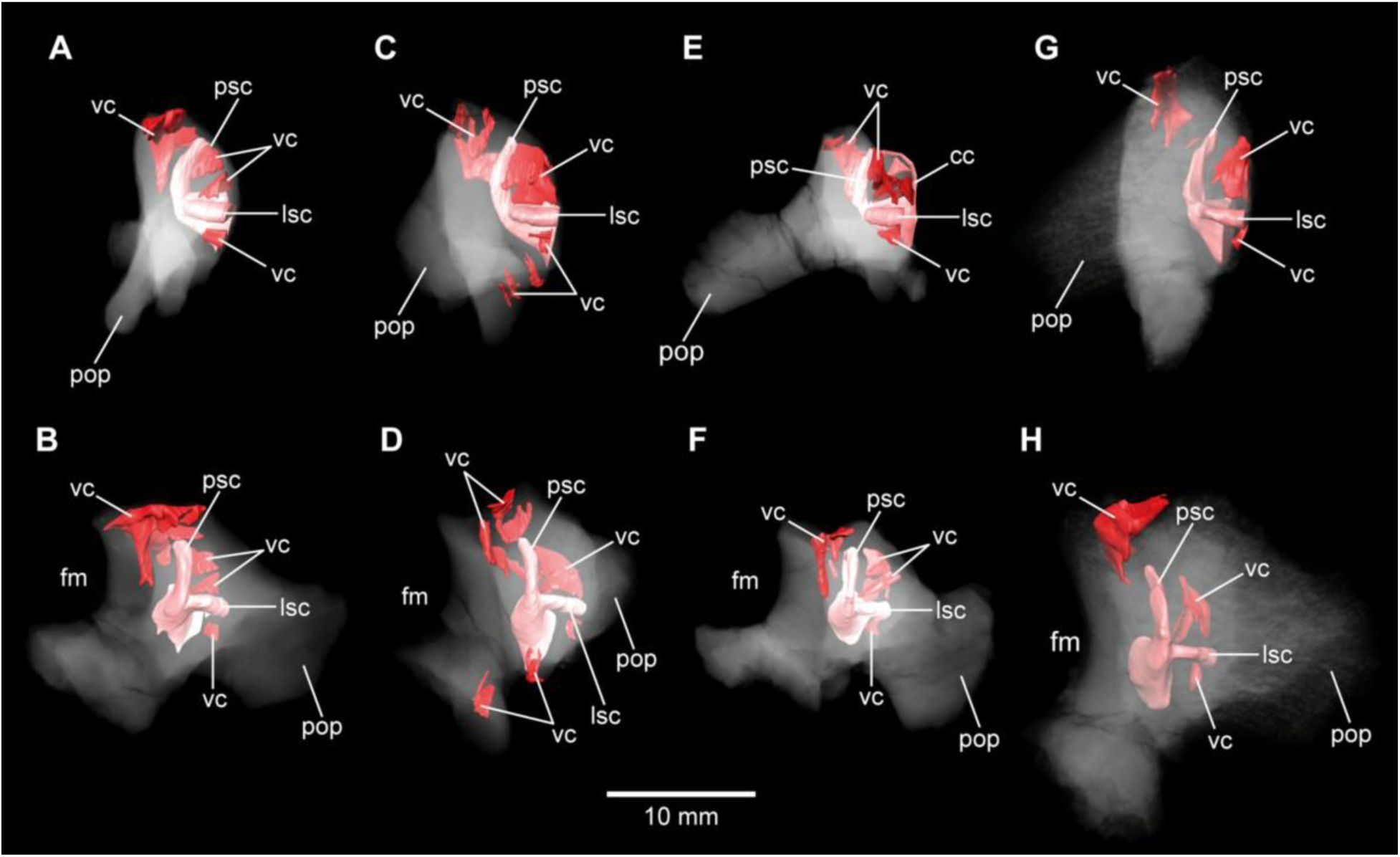
*Europasaurus holgeri*, 3D models of the posterior portions of the endosseous labyrinth in (**A**,**B**) DFMMh/FV 898, (**C**,**D**) DFMMh/FV 981.2, (**E**,**F**) DFMMh/FV 249 and (**G**,**H**) DFMMh/FV 205 in (**A**,**C**,**E**,**G**) anterolateral and (**B**,**D**,**F**,**H**) posterior view. Note that scale mainly applies to posterior perspective (**B**,**D**,**F**,**H**). fm, foramen magnum; lsc, lateral semicircular canal; pop, paroccipital process; psc, posterior semicircular canal; vc, vascular cavity.

The microCT data and our digital reconstructions (<u>Europasaurus holgeri - neuroanatomy - DFMMh/FV - Schade et al. 2023 // MorphoSource</u>) of different *Europasaurus* individuals add to the knowledge of diversity of dinosaur neuroanatomy and allow a better understanding of ontogenetic development. We discuss our findings in context of insights into the lifestyle of this long-necked insular dwarf from the Late Jurassic of Germany.

## Results

### Cranial endocast, innervation and blood supply

As is generally the case in non-maniraptoriform dinosaurs (e.g.^8, 18, 19^), many characteristics of the mid- and hindbrain are not perceivable with certainty on the braincase endocast of DFMMh/FV 581.1 (Fig. 1A), which implies scarce correlation of the actual brain and the inner surface of the endocranial cavity (see^20^ for ontogenetic variations in recent archosaurs).

The endocast suggests low angles in the cerebral and pontine flexures. There is a prominent dorsal expansion, spanning from around the posterodorsal skull roof to approximately the anteroposterior mid-length of the endocast (Figs. 1, 2). In posterior view, the dorsal expansion is T-shaped with a more or less straight top and dorsolateral beams that become dorsoventrally higher and gradually lead over anteriorly to the area where the posterior part of the cerebral hemispheres are expected. In lateral view, the posterior-most extent of the dorsal expansion is separated from the dorsal margin of the medulla oblongata by a concavity.

Anterolaterally to this concavity, the eminence for the vena capitis media is present. Although the respective openings are identifiable on DFMMh/FV 581.1 (close to a kink on the posterodorsal contact between the parietals and the supraoccipital, called ‘external occipital fenestra for the caudal middle cerebral vein‘ in^17^), it was only roughly possible to reconstruct the course of the veins (Figs. 1, 3). There is a large semicircular depression on the posterodorsolateral aspect of the endocast, being anterodorsally bordered by the dorsal expansion and anteroventrally by the eminence of the vena capitis media. On the anterodorsal skull roof, a mediolateral expansion of the endocast possibly marks the position of the cerebral hemisphere (Figs. 2, 3). In lateral view, there is a distinct ventral step on top of the endocast (also present in many other sauropod taxa; see e.g.^7, 21, 22^), between the anterior-most part of the dorsal expansion and the posterior part of the cerebral hemispheres, followed by a slight ascent in anterior direction. The left side of the endocast suggests that the cerebral hemisphere impressions are delimited approximately by the contact between the orbitosphenoid and the laterosphenoid anteriorly, and by the trochlear nerve (CN IV) ventrally. Anteriorly, the orbitosphenoid bears a prominent medial incision for the optic nerve (CN II). Posteroventrally to the optic nerve canal and anteroventrally to the trochlear nerve canal, the canal of the oculomotor nerve (CN III) is situated. On the anteroventral aspect of the endocast, the pituitary reaches about as far ventrally as the ventral-most margin of the medulla oblongata, producing an angle of about 50° to the lateral semicircular canal of the endosseous labyrinth (see^23^). On the anterodorsal aspect of the pituitary, two small and dorsolaterally diverging canals of uncertain identity branch off (Figs. 1–3; Marpmann et al.^17^ labeled the openings as carotid artery: Fig. 13D). In the titanosaur specimen CCMGE 628/12457 and *Sarmientosaurus*, structures of a similar position were identified as sphenoidal arteries^24, 25^. However, in *Bonatitan* and the titanosaur braincase MPCA-PV-80, anterolateral openings on the pituitary, close to the abducens nerve (CN VI) canal, have been assigned to canals leading to the adenohypophysis^7^. Posterolaterally to these canals, the abducens nerve (CN VI) canals trend in an anteroposterior direction (Figs. 1–3; Supplementary Figs. 1–3). The specimen DFMMh/FV 581.1 suggests a natural connection between the pituitary fossa and the left CN VI canal, close to its anterior opening. This condition may be due to breakage, since the microCT data suggests a continuous wall on the right side. In ventrolateral view, the left side of DFMMh/FV 581.1 shows an additional small medial opening dorsally within the depression for CN VI (Figs. 2A,B, 3C,D; Supplementary Figs. 1–3). This opening seems natural, but for preservational reasons, this cannot be confirmed on the right side of the specimen. On the ventrolateral part of the pituitary, two short canals of uncertain identity are branching off ventrolaterally (Figs. 1–3; Supplementary Fig. 1; in *Bonatitan*, anterolateral canals on the ventral portion of the pituitary have been identified as leading to the neurohypophysis^7^). Directly behind, the pituitary bears the long internal carotid canals, branching off ventrolaterally as well.

The endosseous labyrinth is situated within an anteroventrally inclined lateral depression of the endocast, directly ventral to the vena capitis media eminence. Here, an opening is present, leading to the medial aspect of the common crus in DFMMh/FV 581.1 and 1077 (the opening is considerably larger in the latter specimen; Figs. 4, 6; Supplementary Figs. 4, 6).

Whereas the trigeminal (CN V), facial (CN VII) and vestibulocochlear (CN VIII; two openings) nerve canals are mainly anterior to the endosseous labyrinth, the vagal foramen (=jugular foramen for CN IX-XI and jugular vein) and two canals for the hypoglossal nerves are situated posterior to the cochlear duct (Figs. 1–3). Within the depression for CN V, dorsally, a very small opening for the mid cerebral vein is situated on both sides of DFMMh/FV 581.1. However, only the right canal could approximately be reconstructed (Figs. 1A, 2A). Dorsal to the right slit-like opening for CN VII, a small depression is present in DFMMh/FV 581.1.The microCT data do not suggest penetration. Whereas the posterior canals for the hypoglossal nerve (CN XII) are clearly discernable in the microCT data, the anterior ones are not as obvious to detect.

However, because of the expression of their respective openings on the actual fossil, their course could be established (anterior to the proximal openings of the anterior CN XII canals, one depression each is visible on the fossil, however, the microCT data do not suggest a penetration). Marpmann et al.^17^ only identified one hypoglossal canal (CN XII). However, the specimens considered herein, support the presence of two openings on each side.

Anterodorsally to the endosseous labyrinth, the cerebellum seems perceivable as a mediolaterally expanded part of the endocast, almost reaching the trigeminal nerve (CN V) anteriorly and being delimited by the eminence of the vena capitis media posterodorsally (Fig. 1A). Furthermore, a small floccular recess is present close to the mid-length of the anterior semicircular canal in DFMMh/FV 581.1. The ventral aspect of the endocast is anterodorsally inclined and bears a medial ridge (Figs. 1A, 2C, 3A) becoming mediolaterally narrower in anterior direction), reaching between the foramen magnum and the anteroventral portion of the endocast (excluding the pituitary). Posteroventral to the abducens nerve (CN VI), a single median protuberance is present on the endocast, produced by a fossa on the floor of the endocranial cavity (Figs. 2C, 3; Supplementary Figs. 2, 3). In addition, anterodorsally to the proximal openings for the abducens nerve (CN VI), a single median opening is present on the braincase floor, producing a connection to the pituitary fossa (probably for vascularization; see^7, 24^ for arguments on arterial or venous identity). The general osteological configuration of the endocranial floor (Supplementary Figs. 2, 3) seems very similar in the macronarian *Giraffatitan* (^6^: Fig. 117). The anterodorsally incomplete endocranial cavity of DFMMh/FV 581.1, 2 & 3 comprises a volume of about 35 cm^3^ (including the pituitary fossa). A small funnel-like depression anterior to the occipital condyle on the ventral aspect of DFMMh/FV 581.1 ends blindly (Fig. 2D).

#### Endosseous labyrinth

Both vestibular systems are preserved and are ventrally connected to the respective cochlea in DFMMh/FV 581.1 (the semicircular canals of the left inner ear were only vaguely perceptible in some places). Whereas only the left endosseous labyrinth is preserved in DFMMh/FV 1077, only the right one is contained within DFMMh/FV 466+205. The following description is based on the mentioned endosseous labyrinths (Fig. 6). The vertical semicircular canals are relatively long and slender. Dorsoventrally, the anterior semicircular canal reaches considerably higher than the posterior one, and the ASC occupies more of the anteroposterior length of the vestibular system. The common crus is dorsally slightly posteriorly inclined (where preserved). While the PSC forms a low arc, the ASC turns about 180° to contact the common crus dorsomedially. The medial aspect of the common crus is exposed to the endocranial cavity in DFMMh/FV 581.1 and DFMMh/FV 1077 (Figs. 4, 6G; Supplementary Figs. 4, 6). The angle between the ASC and the PSC amounts 80° (measured in dorsal view with the common crus as fixpoint). The lateral semicircular canal is anteroposteriorly short. In dorsal view, its anterior ampulla appears posteriorly shifted, producing a medially concave gap between the ASC and LSC (Fig. 6B,F,J). Such a medial concavity is also present between the LSC and the PSC (best seen in dorsal view). The cochlear duct is approximately as high as the vestibular system dorsoventrally, points anteroventrally and very slightly medially (in DFMMh/FV 581.1 and 1077). In lateral view, the cochlear duct is anteroposteriorly slender with sub-parallel anterior and posterior margins. However, mediolaterally, the cochlear duct is very wide, resulting in an elongated oval-shaped cross section. The fenestra ovalis (Figs. 1A,B, 2A,B, 6; Supplementary Fig. 5) is situated close to the dorsoventral mid-length of the lateral aspect of the cochlear duct (in DFMMh/FV 581.1 and DFMMh/FV 1077).This is also true for the anteroposteriorly oriented fenestra pseudorotunda (Figs. 4, 6D; Supplementary Fig. 4), lying on the posteromedial aspect of the cochlear duct. The hiatus acusticus expresses as an anteromedially open notch (similar to the theropod *Irritator challengeri*^26^) on the actual fenestra pseudorotunda in DFMMh/FV 581.1 (Supplementary Fig. 4).

#### Auditory capabilities

To get a rough idea of the audition of *Europasaurus*, we measured the dorsoventral cochlear duct length of DFMMh/FV 581.1 (c. 16 mm; as outlined by^27^) and the anteroposterior basicranial length (c. 55 mm; from the anterodorsal part of the pituitary fossa to the posterior-most part of the occipital condyle). Based on the equations of^27^, we assume the mean hearing frequency of *Europasaurus* as 2225 Hz and the frequency bandwidth as 3702 Hz (374–4076 Hz).

### Inner ears and cavities of incomplete specimens

In addition to DFMMh/FV 581.1, 2 & 3 (Figs. 1–3, 6A–D; Supplementary Figs. 1–4), eight other braincase specimens (that hold parts of the endosseous labyrinth), assigned to *Europasaurus*, were scanned. DFMMh/FV 1077 (Figs. 4, 6E–H; Supplementary Figs. 5, 6) contains a complete left endosseous labyrinth and was categorized as belonging to an osteological mature individual in Marpmann et al.^17^ (as DFMMh/FV 581.1, 2 & 3). Furthermore, there are two right elements (Figs. 5, 6I–L, 7A,B, 8G,H; Supplementary Figs. 7, 8A,B, 9A, 10G,H, 11E; DFMMh/FV 205, a fragmentary otoccipital, and DFMMh/FV 466, a fragmentary prootic) that were originally found some 10 cm apart from each other in the sedimentary matrix. Whereas DFMMh/FV 205 was thought to belong to a juvenile, DFMMh/FV 466 was supposed to belong to a considerably older individual (both estimations mainly based on size and surface texture^17^). However, DFMMh/FV 205 and DFMMh/FV 466 articulate well with each other and jointly contain most of the endosseous labyrinth and the dorsal portion of the lagena, all in a meaningful manner in respect to size, position and orientation of its compartments. DFMMh/FV 466 is of similar size and texture as the other prootics considered here. DFMMh/FV 205 is considerably smaller than the otoccipitals in the adult specimens. Hence, DFMMh/FV 466+205 are herein interpreted to belong to the same juvenile individual. Furthermore, there are two left fragmentary prootics (Fig. 7C–F; Supplementary Figs. 8C–F, 9B,C; DFMMh/FV 561 and DFMMh/FV 964) containing most of the ASC, the ventral base of the common crus, the anterior ampulla of the LSC and the anterior base of the lagena; both specimens were assigned to relatively mature individuals^17^.

The three right fragmentary otoccipitals DFMMh/FV 249, DFMMh/FV 898 and DFMMh/FV 981.2 (Figs. 8A–F; Supplementary Figs. 10A–F, 11A–D) contain at least the posterior parts of the LSC and the lagena, as well as most of their PSCs; these specimens were assigned to immature individuals^17^.

In general, the morphology of the inner ears contained within these isolated specimens corresponds to what can be observed in DFMMh/FV 581.1 and DFMMh/FV 1077. Since Marpmann et al.^17^ used the vascularization (indicated by surface texture) of *Europasaurus* specimens as a critical character in judging the relative maturity, the inner cavities surrounding the endosseous labyrinths were examined herein.

No discrete cavities could be found in DFMMh/FV 581.1, DFMMh/FV 1077, DFMMh/FV 964 and DFMMh/FV 561 (all considered to represent more or less mature individuals).The otocipitals DFMMh/FV 249, DFMMh/FV 898 and DFMMh/FV 981.2 and the articulated specimens DFMMh/FV 205 (otoccipital) and DFMMh/FV 466 (prootic) show very similar, or corresponding, patterns of inner cavities (Figs. 7, 8; Supplementary Figs. 9, 11). All four otoccipitals show dorsoventrally deep cavities posterodorsal to anteromedial to the PSC, close to the articulation surface with the supraoccipital (except for DFMMh/FV 205, in which this cavity network is not as much extended anteriorly). There are T- (DFMMh/FV 898 and DFMMh/FV 981.2), V- (DFMMh/FV 205), or X- (DFMMh/FV 249) shaped (in cross-section in anterior view), dorsoventrally high and mediolaterally thin structures anterior to the PSC and dorsal to the LSC (close to the articulation surface with the prootic). Additionally, all four otoccipital specimens bear relatively small cavities ventral to the LSC (again, close to the articulation surface with the prootic). DFMMh/FV 981.2 shows dorsoventrally high cavities posteroventrally to the endosseous labyrinth (close to the articulation surface with the basioccipital). Generally, the cavities, likely of vascular purpose, of DFMMh/FV 205 seem not as large and extensive as in the other three otoccipitals. This coincides with their size and assumed relative maturity (DFMMh/FV 205 being the largest, smoothest and, hence, most mature of them^17^). Whereas DFMMh/FV 466 bears a small cavity ventral to the LSC (corresponding to the respective cavity in the otocipital DFMMh/FV 205), no other unequivocal cavities could be found, which is surprising when the V-shaped cavity close to the prootic contact of DFMMh/FV 205 is considered.

## Discussion

### Comparison of neurovascularanatomy and potential ecological implications

Although not as prominent as in *Dicraeosaurus*^6, 28^ and some specimens of *Diplodocus*^29^, the position and morphology of the dorsal expansion of *Europasaurus*, gives a rather ‘upright’ or sigmoidal appearance to the endocast (Fig. 1A; see also^23^). This is partly explained by the (preservational) lack of its olfactory bulb and tract. The first cranial nerve is not expected to be very prominent in many sauropods, especially in the closely-related macronarian taxa *Camarasaurus* and *Giraffatitan*^21, 29^. In contrast, the braincase endocast is rather tubular in some taxa, e.g. the early-diverging sauropodomorph *Buriolestes*^3^, the rebbachisaurid *Nigersaurus taqueti*^30^, and the titanosaur specimen MCCM-HUE-1667^8^. Instead, the endocast of *Europasaurus* seems to be most similar to *Giraffatitan*^6, 21^ (formerly *Brachiosaurus brancai*).

Different from some other sauropod taxa (e.g., *Spinophorosaurus*, *Diplodocus*, *Camarasaurus* and *Sarmientosaurus*; see^25, 29, 31^), there are no discrete canals for vascular features such as e.g., the rostral middle cerebral vein or the orbitocerebral vein on the endocast of *Europasaurus*.

A ventral ridge on the medulla, as seen in *Europasaurus* (Figs. 1A, 2C, 3A), seems to be present, although not as pronounced, in *Thecodontosaurus*^32^, the early-diverging sauropod specimen OUMNH J13596^5^, *Spinophorosaurus*^31^, *Camarasaurus*^29^ and, potentially, *Giraffatitan*^6, 21^ as well.

Although not obvious on the endocast^21^, *Giraffatitan* seems to bear a median fossa posteromedially to the proximal CN VI openings (^6^: Fig. 117); the respective protuberance in *Europasaurus* marks a distinct kink on the endocast (Figs. 2C, 3; Supplementary Figs. 2, 3).

The endocast of *Europasaurus* bears two pairs of canals on the ventrolateral aspect of the pituitary, the posterior of which is interpreted to represent the internal carotid here (Figs. 1A, 2, 3; Supplementary Fig. 1; in accordance to Marpmann et al.^17^: Fig. 13A). Whereas structures identified as the craniopharyngeal canal are present anterior to the carotid artery in the titanosaur specimen CCMGE 628/12457^24^ and the diplodocid specimen MMCh-Pv-232 (assigned to *Leinkupal*^33^), it is situated posteriorly in *Apatosaurus*^34^ (see also^7, 35^ for the – in respect to the internal carotid – anteriorly situated canal of the subcondylar foramen in *Amargasaurus*). However, in these taxa, the craniopharyngeal canal is a singular median canal. This may render the anterior of the two pairs of canals on the ventral aspect of the pituitary in *Europasaurus* the canals for the neurohypophysis^7^. The pituitary of the *Europasaurus* endocast of DFMMh/FV 581.1 does not project much more ventrally than the posteroventral margin of the medulla oblongata. The pituitary is slightly higher dorsoventrally than the anterior semicircular canal (Fig 1A). Usually in sauropods, the pituitary is large and inclined posteroventrally, reaching much more ventrally than the ventral margin of the hindbrain (see e.g. ^21, 25^; see^24^ for an extreme reached in the titanosaur specimen CCMGE 628/12457 with a short anterior semicircular canal and an enormous pituitary). The finding of a relatively small pituitary fossa in *Europasaurus* and early-diverging sauropodomorphs seem to support a close connection of body and pituitary size^3, 36^. The microCT data of DFMMh/FV 581.1 suggest that the right CN VI canal closely passes by the pituitary fossa without a penetration, whereas the left CN VI canal tangents on the pituitary fossa and opens into the latter (Supplementary Fig. 3). The feature of the CN VI canals not penetrating the pituitary seems typical for titanosaurs (e.g.^7, 8, 23^). Whereas Knoll & Schwarz^21^ note such a penetration or connection on the endocast of MB.R.1919, Janensch^6^ originally described penetrating canals in the *Giraffatitan* specimens t 1 and Y 1.

However, the specimen S 66 seems to show CN VI canals rather passing by the pituitary fossa. This may suggest a certain role of individual expressions (*Giraffatitan*), asymmetries (*Europasaurus*) and/or represents a phylogenetically potentially reasonable intermediate state (however, this feature may also be prone to preservational bias).

The endosseous labyrinth of *Europasaurus* (Fig. 6) is most similar to *Giraffatitan*^6^ and *Spinophorosaurus*^31^ in bearing a relatively long ASC and a long lagena. In dorsal view, the anterior ampulla of the short LSC in *Europasaurus* displays a medially concave gap between the ASC and LSC (Fig. 6B,F,J). Similarly, a pronounced concavity is present between the LSC and the PSC (best seen in dorsal view). Both concave gaps (the ASC and the PSC project further laterally than the lateral outline of the LSC reaches medially) are similarly present in many Titanosauriformes (with the exception of FAM 03.064^37^): *Giraffatitan*^6^, *Malawisaurus*^38^, *Sarmientosaurus*^25^, CCMGE 628/12457^24^, *Jainosaurus*^38^, *Ampelosaurus*^22^, *Narambuenatitan*^23^, *Bonatitan*, *Antarctosaurus*, MCF-PVPH 765 and MGPIFD-GR 118^7^, but also in the rebbachisaurids *Limaysaurus* and *Nigersaurus*^39^. In contrast to other sauropods, the anterior portion of the LSC, as well as its lateral-most extent (best seen in dorsal view) seems somewhat posteriorly shifted in the macronarians *Camarasaurus*^29^ and *Europasaurus* (Fig. 6B,F,J; for further discussion see Supplementary Information).

Although the mediolateral width of the lagena seems not associated with auditory capabilities^27^, the lagena of *Europasaurus* is conspicuously thick mediolaterally, especially when compared to its anteroposterior slenderness (Fig. 6). The calculated auditory capacities (based on^27^) impute *Europasaurus* a relatively wide hearing range with a high upper frequency limit (among non-avian dinosaurs^40,41,42^). Walsh et al.^27^ could show a certain correlation between hearing range, complexity of vocalizing and aggregational behaviour in extant reptiles and birds (see also^12, 43^). Following this, it appears plausible that *Europasaurus* lived in groups with conspecifics, which made airborne communication crucial. However, while a given species is likely to perceive sounds within the frequency spectrum it is able to produce, it may be rather unlikely that the full range of frequencies that can be heard is covered by the sound production ability (see^27, 44^). Habitat preferences potentially play a role as well: ‘acoustically cluttered’ habitats like forests seem associated with a tendency towards high frequency intraspecific communication in recent mammals^45^. Together with tropic Late Jurassic conditions in Europe^46^, this may be part of the explanation of the recovered auditory capacities of *Europasaurus*.

### Fragmentary bones and their eco-ontogenetic meaning

An interesting issue are the different morphological ontogenetic stages of DFMMh/FV 466 and DFMMh/FV 205 stated in Marpmann et al.^17^. The authors considered the prootic DFMMh/FV 466 more mature than the otoccipital DFMMh/FV 205. Indeed, DFMMh/FV 466 is about as large as the prootics of DFMMh/FV 581.1, DFMMh/FV 1077, DFMMh/FV 964 and DFMMh/FV 561 (Fig. 7; Supplementary Fig. 8), but the otoccipital DFMMh/FV 205 is much smaller than the ones in DFMMh/FV 581.1 and DFMMh/FV 1077 (and only slightly larger than DFMMh/FV 981.2, DFMMh/FV 898 and DFMMh/FV 249; Fig. 8; Supplementary Fig. 10).

In addition to general size of the specimens, and build and rugosity of articular facets, Marpmann et al.^17^ (see also^47^) defined the morphological ontogenetic stages also by bone surface smoothness, advocating for vascularization: the smoother the surface, the lesser the degree of vascularization and – in tendency – the more mature the individual bone. Our findings support this (Figs. 7, 8; Supplementary Figs. 9, 11). While the bases of individual cavities described herein may represent depressions of articulation areas, their deep penetration into the bone is unambiguous. Apart from this, the described structures might represent sutures.

However, the position and orientation of individual cavities do not conform to what would be expected. Since these cavities make sense in the frame of morphological ontogenetic stages used in Marpmann et al.^17^, they are considered as so far unknown vascular expressions of juvenile *Europasaurus* individuals here.

DFMMh/FV 466 and DFMMh/FV 205 articulate very well with each other, especially on their lateral aspects. Additionally, there are cavities ventral to the LSC that seem to have been continuous originally (Figs. 5, 7A,B, 8G,H; Supplementary Fig. 7). However, whereas the fenestra ovalis is considerably smaller than the vagal foramen in DFMMh/FV 581.1 and DFMMh/FV 1077 (Fig.3B; Supplementary Fig. 5A), it seems that in DFMMh/FV 466+205 this is vice versa (although this impression may be due to the fragmentary nature of the latter two specimens; Fig. 5L; Supplementary Fig. 7F). If DFMMh/FV 466 and DFMMh/FV 205 are in articulation, there is a large gap on their common dorsal aspect (Fig. 5J; Supplementary Fig. 7H). Considering DFMMh/FV 581.1 and DFMMh/FV 1077 and e.g., the braincase of *Massospondylus*^48^, the supraoccipital usually occupies this gap. In case our interpretation of a common bond between DFMMh/FV 466 and DFMMh/FV 205 is misleading and they actually represent two differently-matured individuals, it is still noticeable that the preserved parts of the conjoined endosseous labyrinth of DFMMh/FV 466 and DFMMh/FV 205 display the same features as DFMMh/FV 581.1 and DFMMh/FV 1077 and is anteroposteriorly almost as long as the latter two specimens (Figs. 5, 6; Supplementary Table 1). This suggests an allometric growth between the prootic and otoccipital: during growth, the prootic reaches the ‘adult’ size faster than the otoccipital, producing a surprisingly small paroccipital process (or a surprisingly large prootic) in juvenile individuals of *Europasaurus* (seemingly, also seen in *Massospondylus*^49^), containing a relatively large endosseous labyrinth (see also^50^ for ontogenetic transformations in sauropodomorphs). A relatively large immature endosseous labyrinth is also present in the ornithischians *Dysalotosaurus lettowvobecki*^51^, *Psitaccosaurus lujiatunensis*^13^ and *Triceratops*^52^. Furthermore, the inner ear is relatively large in juveniles of *Massospondylus*^53^ and the extant, precocial ostrich^54^, and stays morphologically relatively stable throughout ontogeny.

The vestibular apparatus detects movements with the aid of endolymphatic fluid and cilia contained within the semicircular canals, which is crucial for locomotion^55^. Thus, a relatively large and morphologically adult-like endosseous labyrinth in expectedly very young individuals of *Europasaurus* suggests that hatchlings had to be light on their feet very fast in this dwarfed sauropod taxon.

## Conclusion

*Europasaurus* has a rather sigmoid general braincase endocast shape, with a comparably large dorsal expansion, two openings for CN XII, an angle of 50° between the pituitary fossa and the LSC, and the ASC is clearly dorsoventrally higher than the PSC (Figs. 1–4, 6). This and additional novel details, such as the highly uniform vascular cavities within the juvenile braincase material (Figs. 7, 8; Supplementary Figs. 9, 11) add to our knowledge about dinosaur neuroanatomy. The relatively small pituitary fossa (Fig. 1A) in an insular dwarf lends support to the idea of being a proxy for body size^36^.

Many sauropods were extremely large land-dwellers as adults, and still, started as tiny hatchlings, indicating enormous growth rates (e.g.^56,57,58^). The threat arising from the discrepancy of several tens of tons between adults and juveniles makes it, among other reasons, unlikely that these animals were able to take good care of their offspring (e.g.^4, 58^).This implies a great mobility early in life (precociality in a broader sense^59^) of sauropods. Although *Europasaurus* represents an island dwarf (adults were probably not as dangerous for their juveniles), having roamed islands about as large as three times nowadays Bavaria, this taxon seemingly retained precociality (and potentially r-strategy^57, 60^) from its large-bodied ancestors. As also suggested by the taphonomic circumstances^14, 16, 17^, (see also Supplementary Information) *Europasaurus* individuals likely stayed in a certain social cohesion, and potentially practiced colonial nesting as is known from other sauropods^2, 60, 61^. In concert with the approximate auditory capabilities offered here, our findings add hints towards the nature of aggregation with a certain complexity of reproductive and social behaviors for these little real-life titans, thriving in Europe some 154 Ma ago.

## Materials and Methods

The articulated braincase specimen of *Europasaurus holgeri*, DFMMh/FV 581.1, together with both loose parietals (FV 581.2 & 3), represents a braincase that is traversed by breakages but not strongly deformed, lacking parts of the anterior and dorsomedial skull roof, as well as the anteromedial walls of the endocranial cavity. The articulated and assembled braincase lacks the frontals, the right orbitosphenoid and laterosphenoid. The parietals are anteriorly, posterodorsomedially and posteriorly incomplete and somewhat deformed (if they fit posteromedially with the supraoccipital they do not fit with the supraoccipital, prootic and laterosphenoid further anteriorly, and vice versa). The braincase of e.g., the macronarian *Giraffatitan brancai* suggests a plain posterior skull roof not exceeding the dorsal extent of the sagittal nuchal crest; this served as an orientation here.

### Macro-photography

All specimens, except DFMMh/FV 581.1, 2 & 3, were documented using a Canon EOS 70D reflex camera equipped with a Canon EFS 10-135 mm objective, extension tubes (13 mm or 21 mm), and a Canon Macro Twin Lite MT-26EX-RT. Light was cross-polarized in order to reduce reflections of the specimen surface. Images were recorded in different focal planes (z-stacks) and subsequently fused with CombineZP (Alan Hadley). All obtained images were optimized for color balance, saturation and sharpness using Adobe Photoshop CS2.

### Micro-computed tomography (microCT)

MicroCT of DFMMh/FV 581.1, 2 & 3 (Figs. 1–3; Supplementary Figs. 1–4) was performed using a Metrotom 1500 (Carl Zeiss Microscopy GmbH, Jena, Germany) in a subsidiary of Zeiss in Essingen. 1804 images were recorded with binning 1 resulting in a DICOM data set (for further details of settings and voxel size see Supplementary Table 1).

All other specimens (Figs. 4, 5, 7, 8; Supplementary Figs. 5–11) were documented with a Xradia MicroXCT-200 (Carl Zeiss Microscopy GmbH, Jena, Germany) of the Imaging center of the Department of Biology, University of Greifswald. 1600 projection images were recorded each, using 0.39x objective lens, with binning 2 (for further details of settings and voxel size for each specimen see Supplementary Table 1). The tomographic images were reconstructed with XMReconstructor software (Carl Zeiss Microscopy GmbH, Jena, Germany), binning 1 (full resolution) resulting in image stacks (TIFF format).

Digital segmentation and measurements were produced utilizing the software Amira (5.6), based on DICOM files (DFMMh/FV 581.1, 2 & 3) and tiff files (remaining material). The microCT data were manually segmented to create 3D surface models. In DFMMh/FV 581.1, 2 & 3, the x- ray absorption of the fossil and the sediment within is quite similar, resulting in low contrast in many places. Furthermore, for preservational reasons (lack of both frontals, right orbitosphenoid, laterosphenoid and loose parietals), the extent of the digital model of the endocast was conservatively estimated on the skull roof and on the anterodorsal region; some asymmetries on the endocast are explained by this circumstance.

## Data availability

The microCT data and neuroanatomical models of all fossil specimens depicted herein are published online, in the repository MorphoSource (<u>Europasaurus holgeri - neuroanatomy - DFMMh/FV - Schade et al. 2023 // MorphoSource</u>)

## Acknowledgements

We are extremely thankful towards Zeiss in Essingen (especially Bastian Zwick and Stephan Tomaschko), the Universitätsmedizin in Greifswald (especially Christopher Nell) for actuating their CT devices for the fossils of *Europasaurus*; further micro-computed tomographies were performed at the Imaging Center of the Department of Biology, University of Greifswald (DFG INST 292/119-1 FUGG; DFG INST 292/120-1 FUGG). We thank Michael ‘Ede’ Kenzler, Jakob Krieger, Georg Brenneis, Jennifer Legat, Steffen Harzsch and Ingelore Hinz-Schallreuter (all University of Greifswald, Germany), together with Benjamin Englich (Dinosaurierpark Münchehagen, Germany) and Serjoscha Evers (University of Fribourg) for their support and discussions. M.S. is supported by the Bogislaw scholarship, University of Greifswald.

## Author contributions

M.S. designed the project, interpreted the data, wrote the manuscript and segmented the CT data of all specimens depicted, whereas C.P. segmented all specimens except for DFMMh/FV 581.1, 2 & 3. N.K. originally unearthed and prepared the fossils. M.S., S.S., C.P. and M.H. prepared the figures. N.K., C.P., M.H. and S.S. contributed to the manuscript.

## Supplementary Information and Figures

### Neurovascularanatomy and potential implications

The fenestra pseudorotunda probably functioned as a pressure relief for the inner ear, and it is considered as the lateral opening of the recessus scalae tympani^1,2,3^. Because a fenestra pseudorotunda seems to be distributed haphazardly among dinosaurs, it was assumed that the three main dinosaur groups (ornithischians, sauropods and theropods) acquired this feature independently^4^. According to the authors, many ornithischians possess a fenestra pseudorotunda (although lost in pachycephalosaurs, ankylosaurs and stegosaurs). Basal sauropodomorphs may originally possessed a fenestra pseudorotunda, whereas derived sauropods lost it (see also^5^). It was suggested that within theropods, a fenestra pseudorotunda did not develop before the origin of coelurosaurs (as well as an additional independent acquisition in abelisauroids^4^). A fenestra pseudorotunda has been identified on the endosseous labyrinth of *Giraffatitan* and the titanosaur MGPIFD-GR 118^6, 7^. In the theropod dinosaur *Irritator*^8^ and in *Europasaurus*, the fenestra pseudorotunda is developed as a mainly anteroposteriorly oriented opening situated medially to the crista interfenestralis (Figs. 4, 6D; Supplementary Fig. 4). In both taxa, the medial wall of the fenestra pseudorotunda is incomplete (possibly from damage, but more likely due to a lack of ossification), producing a hiatus acusticus (Supplementary Fig. 4). This circumstance renders the opening, visible in lateral view of DFMMh/FV 581.1, DFMMh/FV 1077 and DFMMh/FV 466+205 (Figs. 3B, 5L; Supplementary Figs. 5A, 7F), posteriorly to the fenestra ovalis, the vagal foramen for CN IX-XI and the jugular vein. A posterior vagal foramen as in *Irritator* is missing^8^. The unclear pattern of the presence or absence of a fenestra pseudorotunda in dinosaurs may be due to a lack of sufficient fossil preparation or CT data. Alternatively, this aspect has been out of scope in some previous works (these reasons may also hold true for the number of CN XII openings in other taxa). Sobral & Müller^4^ and Sobral et al.^9^ suggest that in taxa without a fenestra pseudorotunda the pressure- relief function has been carried out by the metotic foramen. However, in dinosaurs like the thyreophoran *Struthiosaurus austriacus*, the inner ear seems relatively clear and widely separated from the metotic foramen, which renders a pressure-relief function at least in this taxon rather unlikely (or alternatively, because of other hints towards inferior auditory capacities in this taxon, redundant^10^).

The flocculus is part of the cerebellum and involved in the integration and translation of eye, head and neck movements (VOR, vestibulo-ocular reflex; VCR, vestibulo-collic reflex; see e.g.^11^). In many previous studies, the floccular recess has been treated as a proxy for the actual cerebellar structure, which induced ecological inferences in fossil taxa (e.g.^3, 12, 13^). However, the floccular recess seems not to be a reliable proxy for the size of the actual neural tissue, also containing e.g., vascular tissue^11^, making its potential use problematic^14^. Whereas most sauropod taxa seem not to spot even the slightest floccular recess on the endocast (see e.g.^7, 15^ and references therein), there are some exceptions^16,17,18,19^ including *Giraffatitan*^20^. In case the size of the floccular recess at least partly reflects the actual flocculus in fossil taxa, it potentially holds evidence for superior VOR and VCR in *Europasaurus* and other sauropods that possess this structure (Fig. 1A). Possibly, because of its – for a sauropod – small body size, *Europasaurus* used its neck in a more flexible manner than its large-bodied relatives (see also^5, 21^). Considered in an evolutionary frame, the flocculus seems to leave a perceivable trace on a given endocast rather variably. Seemingly, the more agile early sauropodomorphs still bore relatively large flocculi, which decreased later in size in larger taxa^5, 21^. The presence of a small floccular recess in the secondarily ‘small-bodied’ *Europasaurus* may show a certain evolutionary flexibility in size and, hence, impression on the braincase of the flocculus.

Just as potential implications of the floccular recess for VCR/VOR and agility, the morphology of the semicircular canals in respect to agility or lifestyle (e.g.^22,23,24,25,26^), and the orientation of the lateral semicircular canal in relation to the habitual head posture^27,28,29^, are far from straightforward. If considered as one hint towards a certain orientation of the skull in fossil taxa, the lateral semicircular canal may help to evaluate this issue in concert with aspects such as other osteological correlates and/or ecology (e.g.^10, 29, 30^). If the assumed manner of articulation between the neurocranium DFMMh/FV 581.1 and the rest of the skull is correct^31^ (Fig. 1A,B), the posteroventrally inclining articular facet of the occipital condyle and the horizontal orientation of the lateral semicircular canal of *Europasaurus* suggest an inclined snout of c. 25°, which is similar to some other large-bodied sauropod taxa (e.g.^27, 32,33,34^).

### Geology

Among the most important Mesozoic vertebrate fossil localities in Europe are the marine limestone deposits of the Langenberg quarry near Goslar (Lower Saxony, northern Germany). The quarry on the northern Harz rim belongs to the “Classic Square Mile of Geology”. In this outcrop, evidence of 350 million years of earth history is visible.

The biostratigraphically well dated Jurassic sediments at the Langenberg quarry range from the late Oxfordian to the late Kimmeridgian (see^35^). The predominant lithologies are carbonate beds which are tilted (70 to 80 degrees) to a near-vertical, slightly overturned position during the Harz Mountains orogeny towards the end of the Cretaceous. Sediment composition and invertebrate faunal contents record changes in water depth and salinity, but there is no evidence of subaerial exposure. The limestone deposits are assigned to the Süntel Formation^36^, previously known as the Kimmeridge Formation (German: “Mittlerer Kimmeridge”).

Palaeogeographically, the Langenberg quarry is located in the Lower Saxony Basin which covered most of northern Germany during the Upper Jurassic and Lower Cretaceous and was surrounded by several large paleo-islands. The fossils of *Europasaurus holgeri* are assigned to the lower section of the Upper Kimmeridgian and are 154 million years old.

At least the rock layers 56, 73, and 83^37^ have yielded terrestrial vertebrates that were washed into shallow tidal flats or the sea from a nearby island, most of the other layers contain a partially or purely marine fauna. The finds offer unique insights in the Late Jurassic terrestrial island fauna of northern Germany.

Since 1999, the terrestrial fauna and flora of these paleo-islands have been intensively investigated (see^38^). The quarry is the type locality for the dwarfed basal macronarian sauropod dinosaur *Europasaurus* but also yielded remains of other dinosaurs, such as diplodocid sauropods and stegosaurs. Also, a variety of theropod groups was present including basal Tyrannosauroidea, Allosauroidea, Megalosauroidea cf. *Marshosaurus*, Megalosauridae cf. *Torvosaurus*, and probably Ceratosauria (see^39,40,41^). Non-dinosaurian terrestrial vertebrates include several 3D-preserved remains of pterosaurs, a paramacellodid lizard, and a new genus of atoposaurid crocodilians^42^. Microvertebrate remains recovered by screen-washing are dominated by teeth of fish and crocodyliforms but also included an astonishing range of mammal teeth^43,44,45,46^.

### Taphonomy

The fossils of *Europasaurus* are found only in the bone-rich limestone layer 83, with remarkably few accompanying remains of other animals. With more than 20 individuals discovered so far, it is unlikely that the animals were swimming or that all carcasses were washed from the land into the sea^47, 48^. Instead, it appears likely that the animals lived together in a herd and were located in a tidal plain when they died. This is also supported by the different ontogenetic stages of the discovered *Europasaurus* individuals: from juveniles, subadults to adults. After their sudden death, the *Europasaurus* carcasses were exposed in the shallow water of an intertidal zone for several days. Before their final burial in lime mud, their bodies began to decay. During this decay, the skeletons not only disintegrated but were also chewed upon by small scavengers. Small visible bitemarks on the bone surfaces indicate scavenging by invertebrates, smaller fish, and/or small atoposaurid crocodiles. The absence of theropod bite marks could be an indicator that the sauropod carcasses were deposited in a tidal area inaccessible to large predators.

The excellent preservation of the *Europsaurus* fossils, some still as partly articulated skeletons, as well the presence of at last three articulated sauropod tooth rows indicate very little water movement and little post-mortem transport of the bones before their final embedding in soft mud. Some of the carcasses must have repeatedly lain on top of the tidal banks above the water surface. Exposed to the sun and salty air, the carcasses partially dried out and the spatulate teeth were pulled of the jaw bones by desiccation. The parched and skeletonized cadavers then slowly were covered with sediments.

---

**Supplementary Fig. 1:**
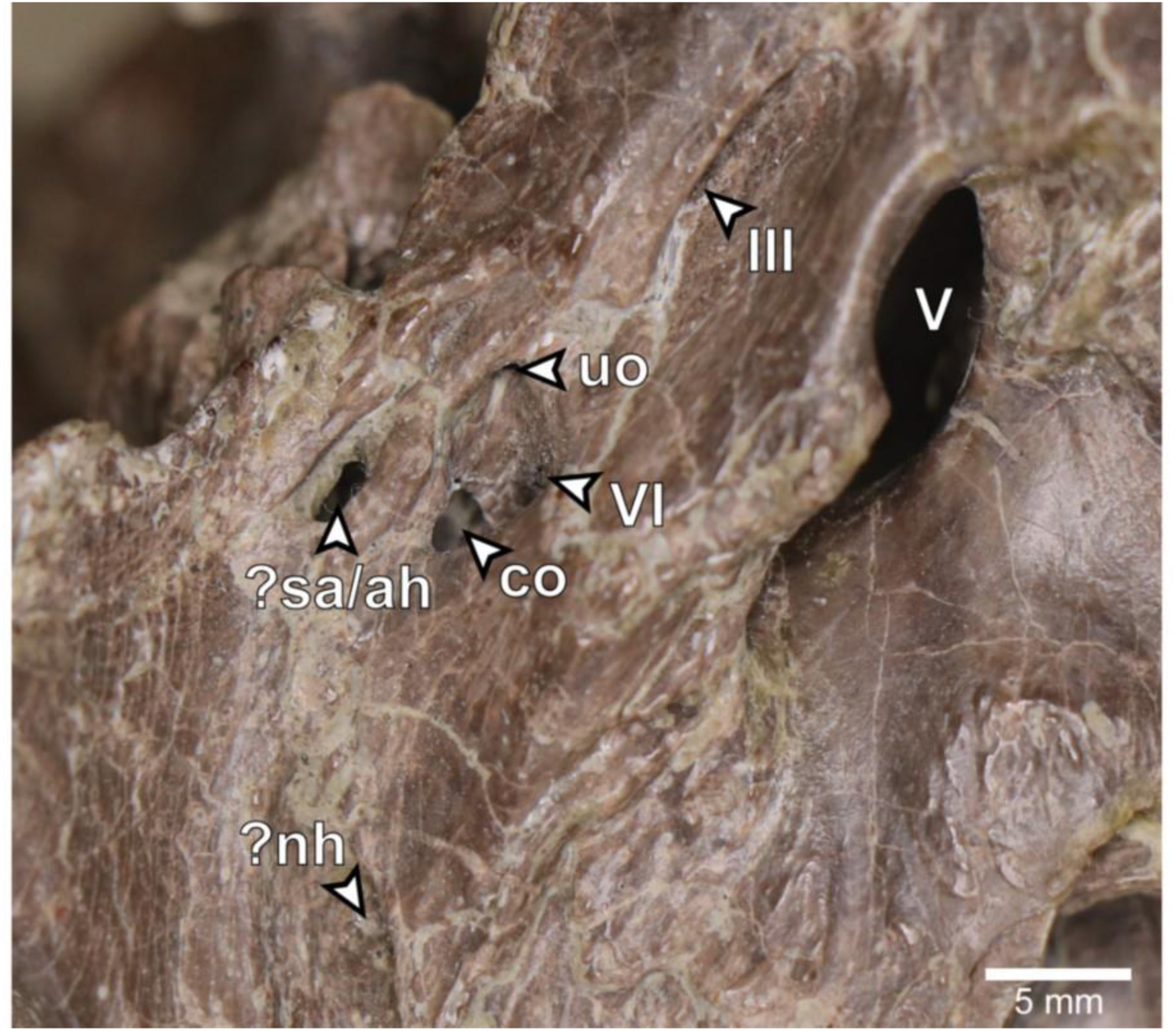
*Europasaurus holgeri*, close-up of left lateral aspect of DFMMh/FV 581.1. ?nh, potential opening for the neurohypophysis; ?sa/ah, potential sphenoidal artery opening/opening for adenohypophysis canal; co, connection between the abducens nerve canal (CN VI) and the pituitary fossa; uo, unclear opening; III, oculomotor nerve opening; V, trigeminal nerve opening; VI, abducens nerve opening.

**Supplementary Fig. 2:**
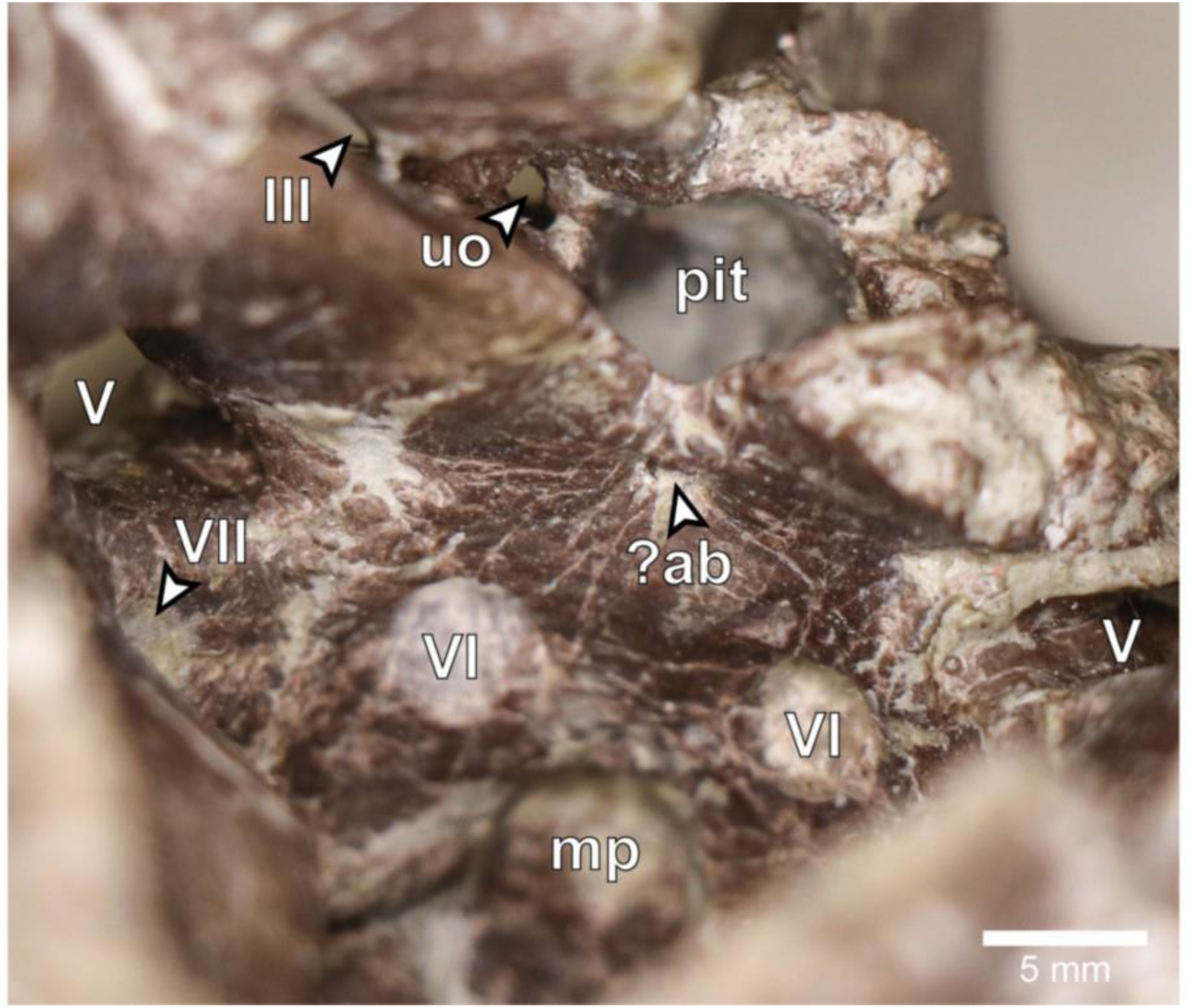
*Europasaurus holgeri*, close-up of anterior endocranial floor of DFMMh/FV 581.1 in dorsal view. ?ab, potential basilar artery opening; mp, median fossa producing the median protuberance on the ventral braincase endocast; pit, pituitary fossa; uo, unclear opening; III, oculomotor nerve opening; V, trigeminal nerve opening; VI, abducens nerve opening; VII, facial nerve opening.

**Supplementary Fig. 3:**
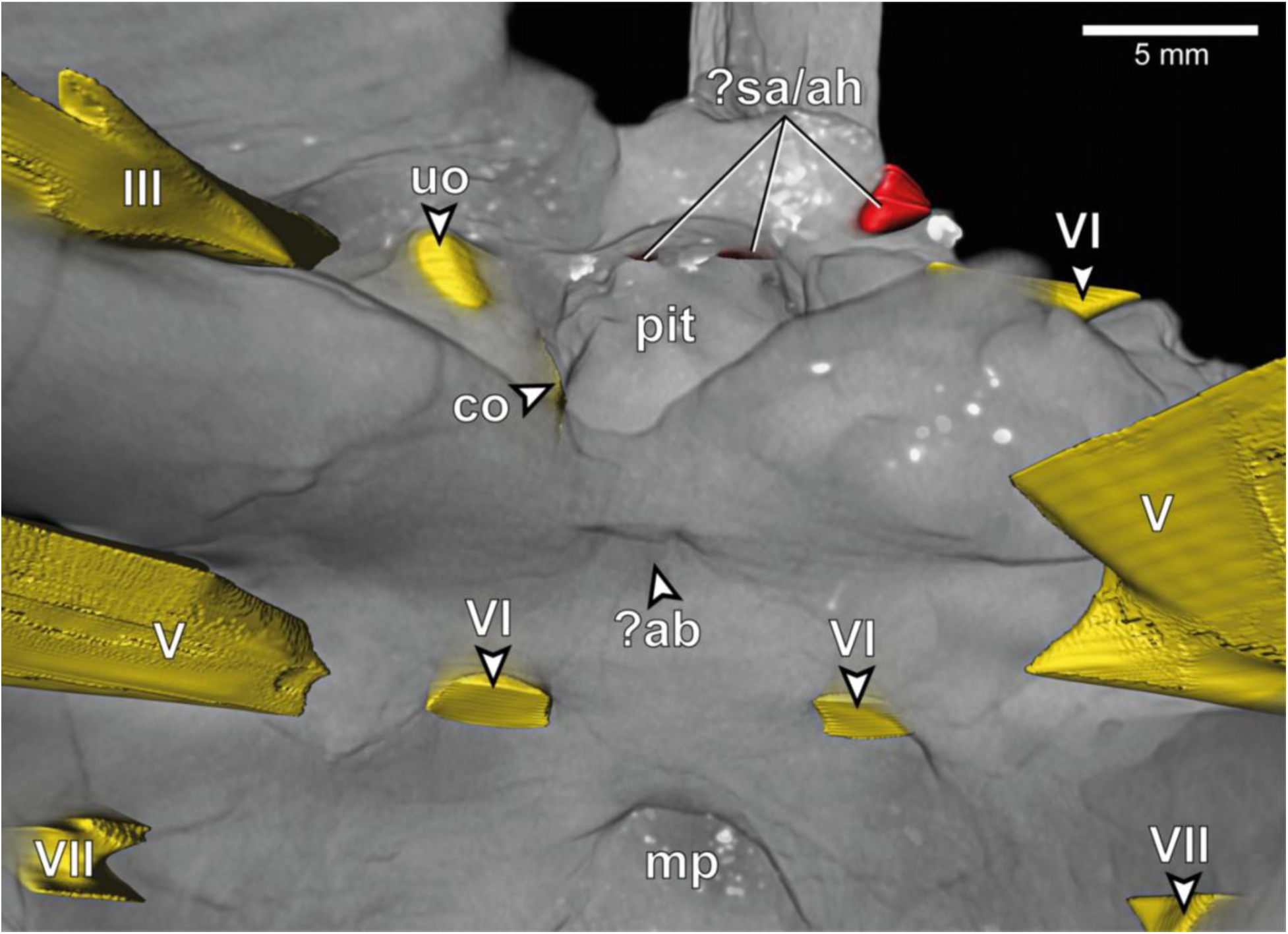
*Europasaurus holgeri*, close-up of 3D model of anterior endocranial floor of DFMMh/FV 581.1 in dorsal view. ?ab, potential basilar artery opening; ?sa/ah, potential sphenoidal artery opening/opening for adenohypophysis canal; co, connection between the abducens nerve canal (CN VI) and the pituitary fossa; mp, median fossa producing the median protuberance on the ventral braincase endocast; uo, unclear opening; pit, pituitary fossa; III, oculomotor nerve opening; V, trigeminal nerve opening; VI, abducens nerve opening; VII, facial nerve opening.

**Supplementary Fig. 4:**
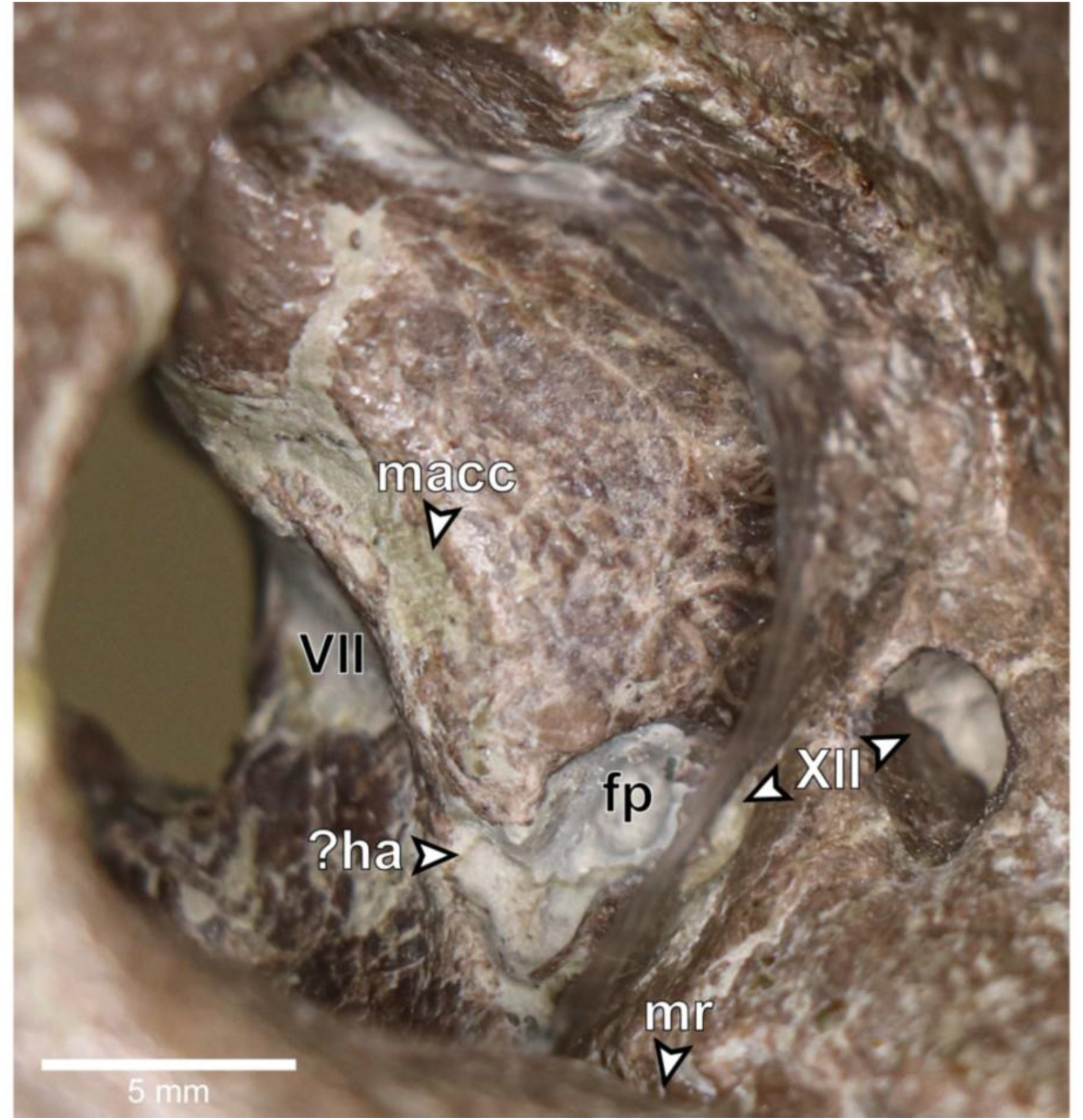
*Europasaurus holgeri*, close-up of right posterior endocranial wall of DFMMh/FV 581.1, viewed through the foramen magnum. ?ha, potential hiatus acusticus; fp, fenestra pseudorotunda; macc, medial aspect of common crus; mr, medial trough producing the medial ridge on the ventral braincase endocast; VII, facial nerve opening; XII, hypoglossal nerve openings.

**Supplementary Fig. 5:**
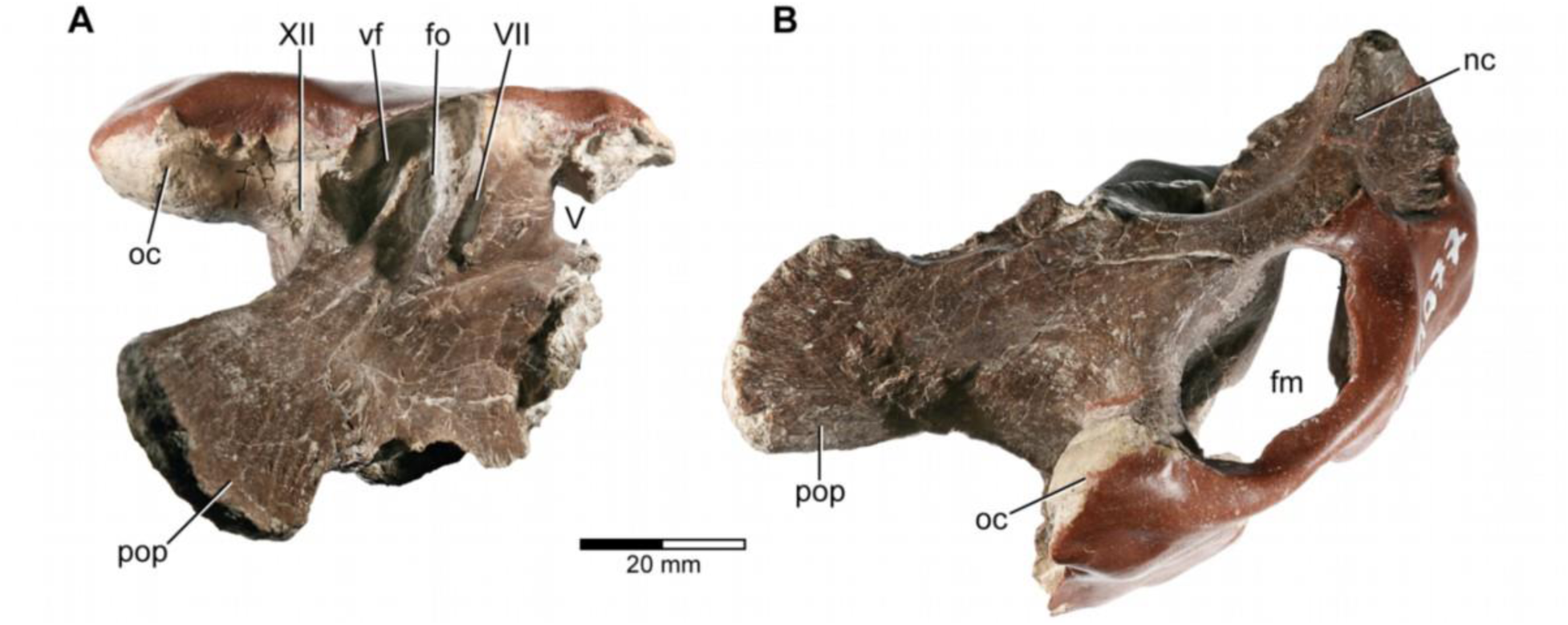
*Europasaurus holgeri*, fragmentary braincase DFMMh/FV 1077 in (**A**) ventral and (**B**) posterior view. fm, foramen magnum; fo, fenestra ovalis; nc, nuchal crest; oc, occipital condyle; pop, paroccipital process; vf, vagal foramen; V, trigeminal nerve opening; VII, facial nerve opening; XII, hypoglossal nerve opening.

**Supplementary Fig. 6:**
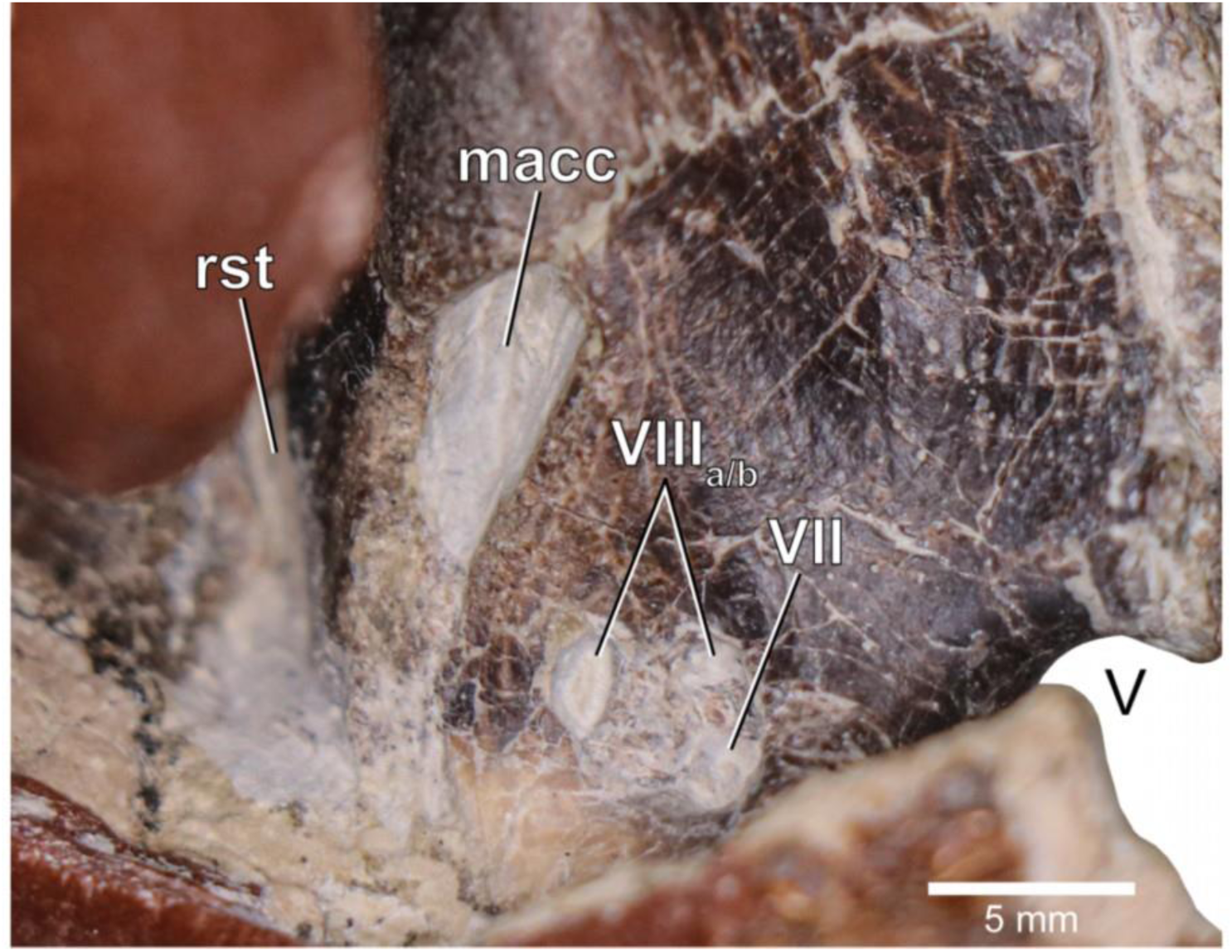
*Europasaurus holgeri*, close-up of medial aspect of the fragmentary braincase DFMMh/FV 1077. macc, medial aspect of common crus; rst, recessus scalae tympani; V, trigeminal nerve opening; VII, facial nerve opening; VIIIa/b, both openings of the vestibulocochlear nerve.

**Supplementary Fig. 7:**
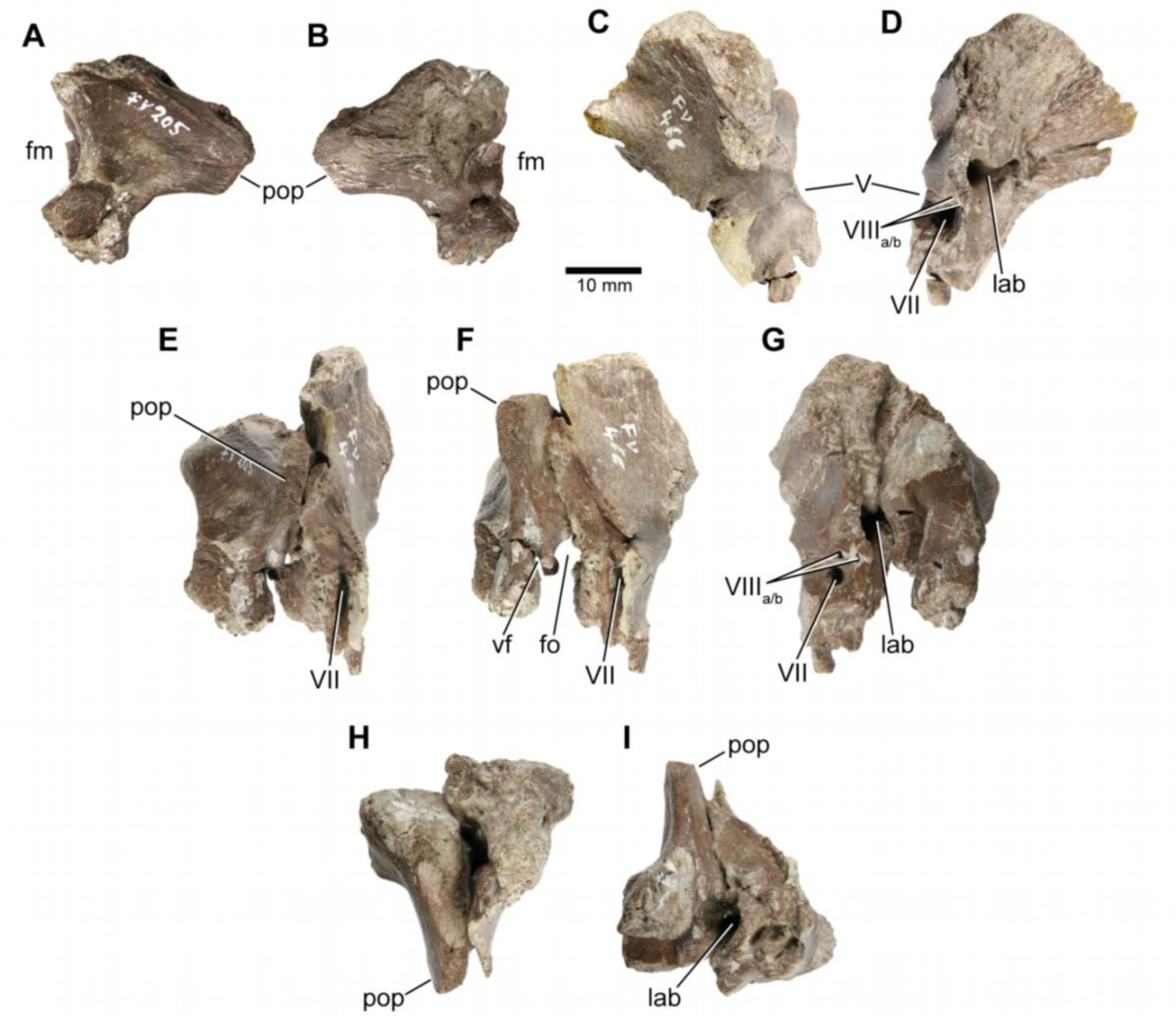
*Europasaurus holgeri*, isolated otoccipital (DFMMh/FV 205; **A**,**B**) and prootic (DFMMh/FV 466; **C**,**D**) in (**A**) posterior, (**B**) anterior, (**C**) lateral and (**D**) medial view; prootic and otoccipital conjoined in (**E**) posterolateral, (**F**) lateral, (**G**) medial, (**H**) dorsal and (**I**) ventral view. Note that scale mainly applies to **A** and **B**. fm, foramen magnum; fo, fenestra ovalis; lab, endosseous labyrinth; pop, paroccipital process; vf, vagal foramen; V, trigeminal nerve opening; VII, facial nerve opening; VIIIa/b, both openings of the vestibulocochlear nerve.

**Supplementary Fig. 8:**
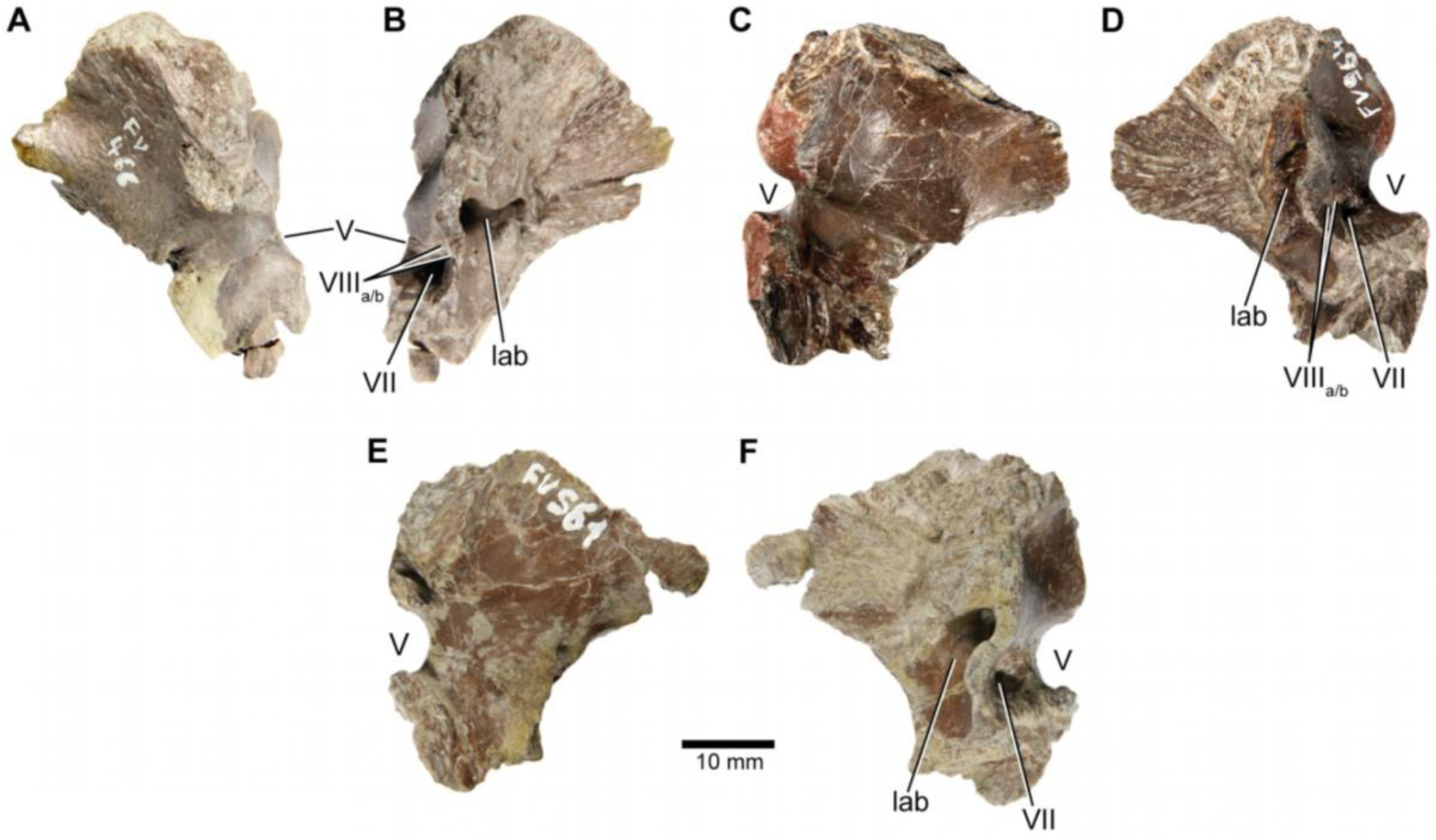
*Europasaurus holgeri*, isolated prootics (DFMMh/FV 466, **A**,**B**; DFMMh/FV 964, **C**,**D**; DFMMh/FV 561, **E**,**F**) in (**A**,**C**,**E**) lateral and (**B**,**D**,**F**) medial view. Note that scale mainly applies to lateral perspective (**A**,**C**,**E**). lab, endosseous labyrinth; V, trigeminal nerve opening; VII, facial nerve opening; VIIIa/b, both openings of the vestibulocochlear nerve.

**Supplementary Fig. 9:**
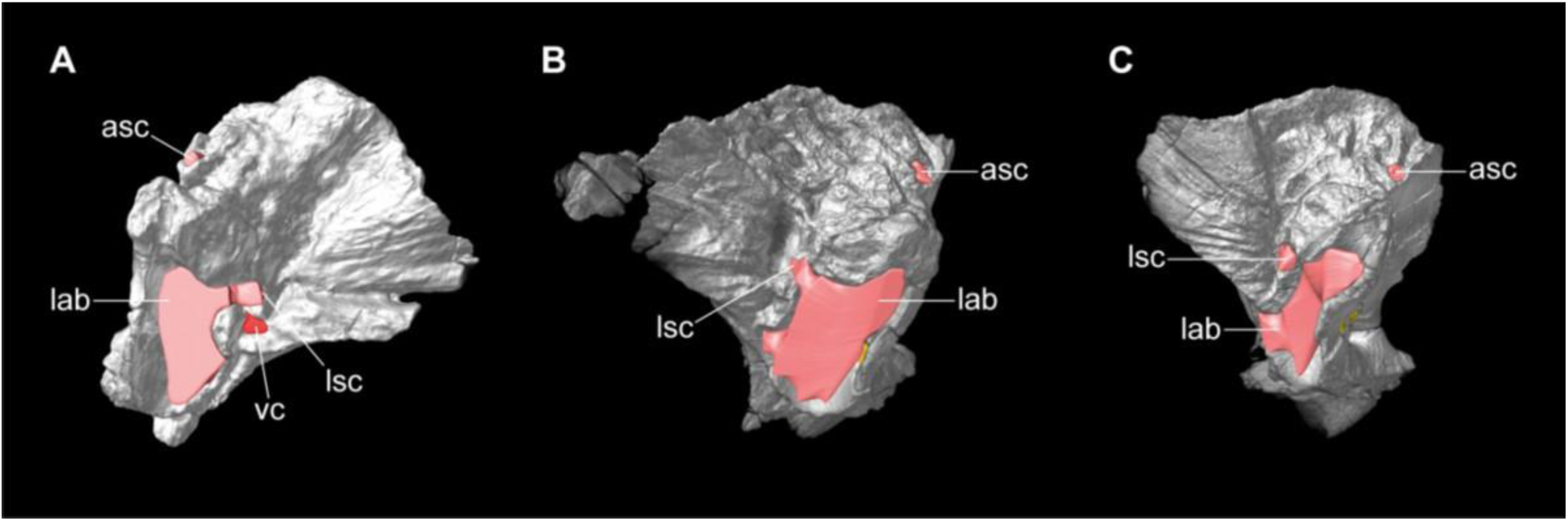
*Europasaurus holgeri*, 3D models of isolated prootics and inner features (DFMMh/FV 466, **A**; DFMMh/FV 561, **B**; DFMMh/FV 964, **C**) in (**A**-**C**) medial view. Note that models are not scaled. asc, anterior semicircular canal; lab, endosseous labyrinth; lsc, lateral semicircular canal; vc, vascular cavity.

**Supplementary Fig. 10:**
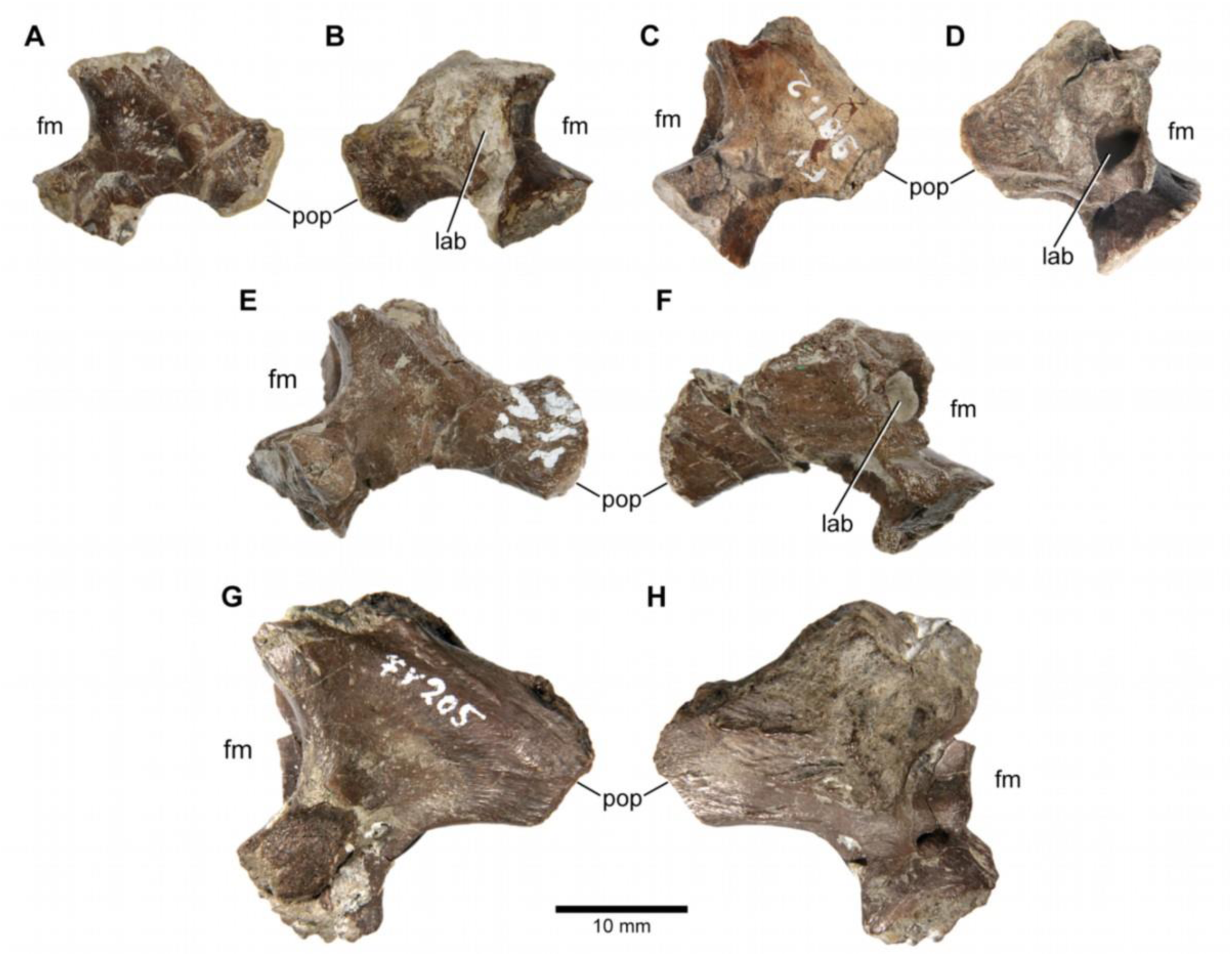
*Europasaurus holgeri*, isolated otoccipitals (DFMMh/FV 898, **A**,**B**; DFMMh/FV 981.2, **C**,**D**; DFMMh/FV 249, **E**,**F**; DFMMh/FV 205, **G**,**H**) in (**A**,**C**,**E**,**G**) posterior and (**B**,**D**,**F**,**H**) anterior view. Note that scale mainly applies to posterior perspective (**A**,**C**,**E**,**G**). fm, foramen magnum; lab, endosseous labyrinth; pop, paroccipital process.

**Supplementary Fig. 11:**
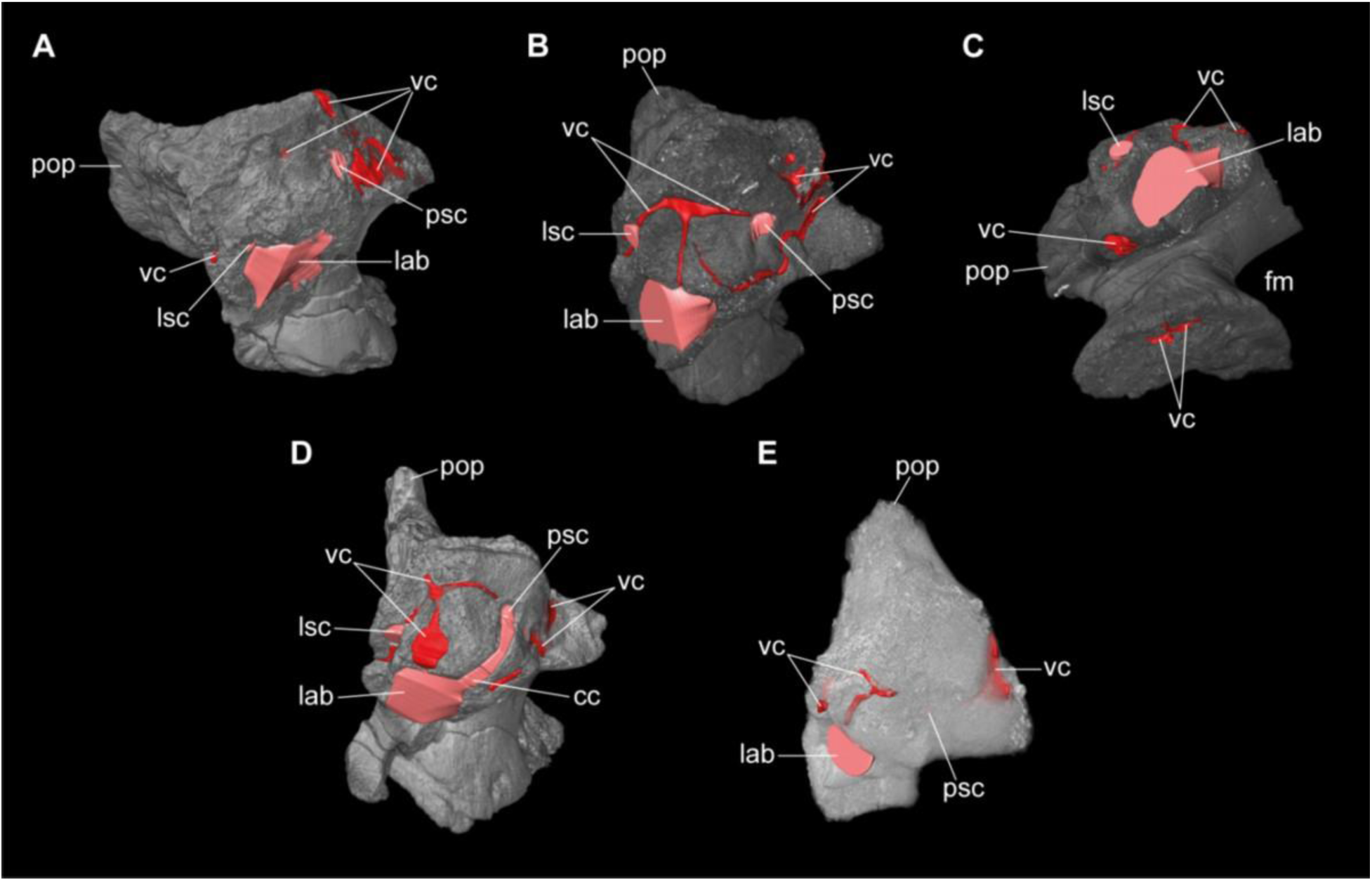
*Europasaurus holgeri*, 3D models of isolated otoccipitals and inner features (DFMMh/FV 898, **A**; DFMMh/FV 981.2, **B**,**C**; DFMMh/FV 249, **D**; DFMMh/FV 205, **E**) in (**A**,**B**,**D**,**E**) anterodorsomedial and (**C**) ventral view. Note that models are not scaled. cc, common crus; fm, foramen magnum; lab, endosseous labyrinth; lsc, lateral semicircular canal; pop, paroccipital process; psc, posterior semicircular canal; vc, vascular cavity.

## References

1. Bates, K. et al. Temporal and phylogenetic evolution of the sauropod dinosaur body plan. R. Soc. Open Sci. 3, 10.1098/rsos.150636 (2016).

2. Pol, D., Otero, A., Apaldetti, C. & Martínez, R. N. Triassic sauropodomorph dinosaurs from South America: The origin and diversification of dinosaur dominated herbivorous faunas. J. S. Am. Earth Sci. 107, 103145, 10.1016/j.jsames.2020.103145 (2021).

3. Müller, R. T., Ferreira, J. D., Pretto, F. A., Bronzati, M. & Kerber L. The endocranial anatomy of *Buriolestes schultzi* (Dinosauria: Saurischia) and the early evolution of brain tissues in sauropodomorph dinosaurs. J. Anat. 238, 809–827, https://doi.org/10.1111/joa.13350 (2021).

4. Sander, P. M., et al. Biology of the sauropod dinosaurs: the evolution of gigantism. Biol. Rev. 86, 117–155, https://doi.org/10.1111/j.1469-185X.2010.00137.x (2010).

5. Bronzati, M., Benson, R. B. J. & Rauhut, O. W. M. Rapid transformation in the braincase of sauropod dinosaurs: integrated evolution of the braincase and neck in early sauropods? Palaeontology 61, 289–302, 10.1111/pala.12344 (2018).

6. Janensch, W. Die Schädel der Sauropoden *Brachiosaurus*, *Barosaurus* und *Dicraeosaurus* aus den Tendaguru-Schichten Deutsch-Ostafrikas. *Palaeontographica*, Suppl. 7, 1 (2), 147–298. (1935).

7. Paulina-Carabajal, A. Neuroanatomy of titanosaurid dinosaurs from the Upper Cretaceous of Patagonia, with comments on endocranial variability within sauropoda. Anat. Rec. (Hoboken, N.J.: 2007) 295, 2141–2156, 10.1002/ar.22572 (2012).

8. Knoll, F., Witmer, L. M., Ridgely, R. C., Ortega, F., Sanz, J. L. A new titanosaurian braincase from the Cretaceous “Lo Hueco” locality in Spain sheds light on neuroanatomical evolution within titanosauria. PLoS ONE 10(10): e0138233, doi:10.1371/journal.pone.0138233 (2015).

9. Schwab, J. A., Young, M. T., Neenan, J. M., Brusatte, S. L. Inner ear sensory system changes as extinct crocodylomorphs transitioned from land to water. Proc. Natl. Acad. Sci. USA. 117, 10422–10428, https://doi.org/10.1073/pnas.2002146117 (2020).

10. Schwab, J. A., Young, M. T., Herrera, Y., Witmer, L. M., Walsh, S. A., Katsamenis, O. L. & Brusatte, S. L. The braincase and inner ear of *‘Metriorhynchus’* cf. ‘M.’ *brachyrhynchus* – implications for aquatic sensory adaptations in crocodylomorphs, J. Vertebr. Paleontol. DOI: 10.1080/02724634.2021.1912062 (2021).

11. Ezcurra, M. D. et al. Enigmatic dinosaur precursors bridge the gap to the origin of Pterosauria. Nature 588 (7838), 445–449, https://doi.org/10.1038/s41586-020-3011-4 (2020).

12. Hanson, M., Hoffman, E. A., Norell, M. A. & Bhullar, B. S. The early origin of a birdlike inner ear and the evolution of dinosaurian movement and vocalization. Science 372 (6542), 601–609, https://doi.org/10.1126/science.abb4305 (2021).

13. Bullar, C. M., Zhao, Q., Benton, M. J., Ryan, M. J. Ontogenetic braincase development in *Psittacosaurus lujiatunensis* (Dinosauria: Ceratopsia) using micro-computed tomography. PeerJ 7:e7217, DOI 10.7717/peerj.7217 (2019).

14. Sander, P. M., Mateus, O., Laven, T. & Knötschke, N. Bone histology indicates insular dwarfism in a new Late Jurassic sauropod dinosaur. Nature 441, 739–741 (2006).

15. Stein, K., Csiki, Z., Rogers, K. C., Weishampel, D. B., Redelstorff, R., Carballido, J. L. & Sander, P. M. Small body size and extreme cortical bone remodeling indicate phyletic dwarfism in *Magyarosaurus dacus* (Sauropoda: Titanosauria). PNAS USA 107, 9258–9263 (2010).

16. Carballido, J. L. & Sander, M. P. Postcranial axial skeleton of *Europasaurus holgeri* (Dinosauria, Sauropoda) from the Upper Jurassic of Germany: implications for sauropod ontogeny and phylogenetic relationships of basal Macronaria. J. Syst. Palaeontol. 3, 335–387 (2014).

17. Marpmann, J. S., Carballido, J. L., Sander, M. P. & Knötschke, N. Cranial anatomy of the Late Jurassic dwarf sauropod *Europasaurus holgeri* (Dinosauria, Camarasauromorpha): ontogenetic changes and size dimorphism, J. Syst. Palaeontol. DOI: 10.1080/14772019.2013.875074 (2014).

18. Witmer, L. M. & Ridgely, R. C. New insights into the brain, braincase, and ear region of tyrannosaurs (Dinosauria, Theropoda), with implications for sensory organization and behavior. Anat. Rec. 292(9), 1266–1296, https://doi.org/10.1002/ar.20983 (2009).

19. Becerra, M., Paulina-Carabajal, A., Cruzado-Caballero, P. & Taborda, J. First endocranial description of a South American hadrosaurid: The neuroanatomy of *Secernosaurus koerneri* from the Late Cretaceous of Argentina. APP 63; 10.4202/app.00526.2018 (2018).

20. Watanabe, A. et al. Are endocasts good proxies for brain size and shape in archosaurs throughout ontogeny? J. Anat. 234, 291–305; 10.1111/joa.12918 (2019).

21. Knoll, F. & Schwarz-Wings, D. Palaeoneuroanatomy of *Brachiosaurus*. Ann. Paléontol. 95, 165–175; 10.1016/j.annpal.2009.06.001 (2009).

22. Knoll, F., Ridgely, R. C., Ortega, F., Sanz, J. L. & Witmer, L. M. Neurocranial osteology and neuroanatomy of a late Cretaceous titanosaurian sauropod from Spain (*Ampelosaurus* sp.). PloS ONE 8, e54991; 10.1371/journal.pone.0054991 (2013).

23. Paulina-Carabajal, A., Filippi, L., & Knoll, F. Neuroanatomy of the titanosaurian sauropod *Narambuenatitan palomoi* from the Upper Cretaceous of Patagonia, Argentina. Pe APA 20(2): 1–9 (2020).

24. Sues, H.-D., Averianov, A., Ridgely, R. C. & Witmer, L.M. Titanosauria (Dinosauria, Sauropoda) from the Upper Cretaceous (Turonian) Bissekty Formation of Uzbekistan. J. Vertebr. Paleontol. DOI: 10.1080/02724634.2014.889145 (2015).

25. Martínez, R. D. F., Lamanna, M. C., Novas, F. E., Ridgely, R. C., Casal, G. A., Martínez, J. E., Vita, J. R. & Witmer, L. M. A basal lithostrotian titanosaur (Dinosauria: Sauropoda) with a complete skull: implications for the evolution and paleobiology of Titanosauria. PLoS ONE, 11(4), e0151661 (2016).

26. Schade, M., Rauhut, O. W. M. & Evers, S. W. Project: Schade, et al. *Irritator challengeri* SMNS 58022 neuroanatomy. MorphoSource, available at https://www.morphosource.org/Detail/ProjectDetail/Show/project_id/951 (2020).

27. Walsh, S. A., Barrett, P. M., Milner, A. C., Manley, G. & Witmer, L. M. Inner ear anatomy is a proxy for deducing auditory capability and behaviour in reptiles and birds. Proc. R. Soc. B 276, 1355–1360 (2009).

28. Paulina-Carabajal, A., Coria, R. A., Currie, P. J. & Koppelhus, E. B. A natural cranial endocast with possible dicraeosaurid (Sauropoda, Diplodocoidea) affinities from the Lower Cretaceous of Patagonia. Cretac. Res. 84, 437–441, 10.1016/j.cretres.2017.12.001 (2017).

29. Witmer, L. M., Ridgely, R. C., Dufeau, D. L. & Semones, M. C. Using CT to peer into the past: 3D visualization of the brain and ear regions of birds, crocodiles, and nonavian dinosaurs. In Anatomical imaging: towards a new morphology (eds Endo, H. & Frey, R.) 67–87 (Springer, 2008).

30. Sereno, P.C. et al. Structural extremes in a Cretaceous dinosaur. PLoS ONE 2(11): e1230. doi:10.1371/journal.pone.0001230 (2007).

31. Knoll, F., Witmer, L. M., Ortega, F., Ridgely, R. C. & Schwarz-Wings, D. The braincase of the basal sauropod dinosaur *Spinophorosaurus* and 3D reconstructions of the cranial endocast and inner ear. PLoS ONE 7(1): e30060, doi:10.1371/journal.pone.0030060 (2012).

32. Ballell, A., King, L., Neenan, J., Rayfield, E. & Benton, M. The braincase, brain and palaeobiology of the basal sauropodomorph dinosaur *Thecodontosaurus antiquus*. Zool. J. Linn. Soc. 193, 10.1093/zoolinnean/zlaa157 (2020).

33. Garderes, J.P., Gallina, P.A., Whitlock, J.A. & Toledo, N., Neuroanatomy of a diplodocid sauropod dinosaur from the Lower Cretaceous of Patagonia, Argentina, Cretac. Res. https://doi.org/10.1016/j.cretres.2021.105024 (2022).

34. Balanoff, A. M., Bever, G. S. & Ikejiri, T. The braincase of *Apatosaurus* (Dinosauria: Sauropoda) based on computed tomography of a new specimen with comments on variation and evolution in sauropod neuroanatomy. Am. Mus. Novit. 3677, 1–32; 10.1206/591.1 (2010).

35. Paulina-Carabajal, A., Carballido, J. L. & Currie, P. J. Braincase, neuroanatomy, and neck posture of *Amargasaurus cazaui* (Sauropoda, Dicraeosauridae) and its implications for understanding head posture in sauropods. J. Vertebr. Paleontol. 34, 870–882, 10.1080/02724634.2014.838174 (2014).

36. Edinger, T. The pituitary body in giant animals fossil and living: a survey and a suggestion. Q. Rev. Biol. 17, 31–45 (1942).

37. Knoll, F., Lautenschlager, S., Valentin, X., Díez Díaz, V., Pereda Suberbiola, X. & Garcia, G. First palaeoneurological study of a sauropod dinosaur from France and its phylogenetic significance. PeerJ 7:e7991 DOI 10.7717/peerj.7991 (2019).

38. Andrzejewski, K. A., Polcyn, M. J., Winkler, D. A., Gomani Chindebvu, E. & Jacobs, L.L. The braincase of *Malawisaurus dixeyi* (Sauropoda: Titanosauria): A 3D reconstruction of the brain endocast and inner ear. PLoS ONE 14(2): e0211423, https://doi.org/10.1371/journal.pone.0211423 (2019).

39. Paulina-Carabajal, A. & Calvo, J. O. Re-description of the braincase of the rebbachisaurid sauropod *Limaysaurus tessonei* and novel endocranial information based on CT scans. An. Acad. Bras. Cienc. 93: e20200762, DOI. 10.1590/0001-3765202120200762 (2021).

40. Lautenschlager, S., Rayfield, E.J., Altangerel, P., Zanno, L. E., Witmer, L. M. The endocranial anatomy of therizinosauria and its implications for sensory and cognitive function. PLoS ONE 7(12): e52289, doi:10.1371/journal.pone.0052289 (2012).

41. King, J. L., Sipla, J. S., Georgi, J. A., Balanoff, A. M. & Neenan, J. M. The endocranium and trophic ecology of *Velociraptor mongoliensis*. J. Anat. 237: 861–869 (2020).

42. Sakagami, R., Kawabe, S. Endocranial anatomy of the ceratopsid dinosaur *Triceratops* and interpretations of sensory and motor function. PeerJ 8:e9888, http://doi.org/10.7717/peerj.9888 (2020).

43. Gleich, O., Dooling, R. J. & Manley, G. A. Audiogram, body mass, and basilar papilla length: correlations in birds and predictions for extinct archosaurs. Naturwissenschaften 92, 595–589, doi:10.1007/s00114-005-0050-5 (2005).

44. Senter, P. Voices of the past: a review of Paleozoic and Mesozoic animal sounds. Hist. Biol. 20:4, 255–287, DOI: 10.1080/08912960903033327 (2008).

45. Charlton, B. D., Owen, M. A. & Swaisgood, R. R. Coevolution of vocal signal characteristics and hearing sensitivity in forest mammals. Nat. Commun. 10, 2778, https://doi.org/10.1038/s41467-019-10768-y (2019).

46. Armstrong, H. A. et al. Hadley circulation and precipitation changes controlling black shale deposition in the Late Jurassic Boreal Seaway. Paleoceanography 31, 1041–1053, doi:10.1002/2015PA002911 (2016).

47. Benton, M. J. et al. Dinosaurs and the island rule: the dwarfed dinosaurs from Hateg Island. Palaeogeogr. Palaeoclimatol. Palaeoecol. 293, 438–454 (2010).

48. Chapelle, K. E. J. & Choiniere, J. N. A revised cranial description of *Massospondylus carinatus* Owen (Dinosauria: Sauropodomorpha) based on computed tomographic scans and a review of cranial characters for basal Sauropodomorpha. PeerJ 6, e4224, 10.7717/peerj.4224 (2018).

49. Sues, H.-D., Reisz, R. R., Hinic, S. & Raath, M.A. On the skull of *Massospondylus carinatus* Owen, 1854 (Dinosauria: Sauropodomorpha) from the Elliot and Clarens formations (Lower Jurassic) of South Africa. Ann. Carnegie Mus. 73, 239–257 (2004).

50. Fabbri, M. et al. A shift in ontogenetic timing produced the unique sauropod skull. Evol.; int. j. org. evol. 75, 819–831, 10.1111/evo.14190 (2021).

51. Lautenschlager, S. & Hübner, T. Ontogenetic trajectories in the ornithischian endocranium. J. Evol. Biol. 26, 2044–2050, 10.1111/jeb.12181 (2013).

52. Morhardt, A. C., et al. Study of endocranial & ontogeny in the Late Cretaceous non-avian dinosaur genus *Triceratops* using computed tomography & 3D visualization. Poster (2018).

53. Neenan, J. M., Chapelle, K. E. J., Fernandez, V. & Choiniere, J. N. Ontogeny of the *Massospondylus* labyrinth: implications for locomotory shifts in a basal sauropodomorph dinosaur. Palaeontology 62, 255–265, 10.1111/pala.12400 (2019).

54. Romick, C. A. Ontogeny of the brain endocasts of Ostriches (Aves: Struthio camelus) with implications for interpreting extinct dinosaur endocasts [Undergraduate thesis, Ohio University]. OhioLINK Electronic Theses and Dissertations Center. http://rave.ohiolink.edu/etdc/view?acc_num=ouashonors1368018907 (2013).

55. Benson, R. B. J., Starmer-Jones, E., Close, R. A. & Walsh, S. A. Comparative analysis of vestibular ecomorphology in birds. J. Anat. 231, 990–1018, https://doi.org/10.1111/joa.12726 (2017).

56. Carpenter, K. Eggs, nests and baby dinosaurs: A look at dinosaur reproduction. 336p (Indiana University Press, 1999).

57. Hallett, M. & Wedel, M. J. The sauropod dinosaurs. Life in the age of giants. 320p (Johns Hopkins University Press, 2016).

58. Curry Rogers, K., Whitney, M., D’Emic, M. & Bagley, B. Precocity in a tiny titanosaur from the Cretaceous of Madagascar. Science 352, 450–453, 10.1126/science.aaf1509 (2016).

59. Dial, K.P. Evolution of avian locomotion: correlates of flight style, locomotor modules, nesting biology, body size, development, and the origin of flapping flight. Ornithology 120(4), 941–952 (2003).

60. Sander, P. M., Peitz, C., Jackson, F. D. & Chiappe, L. M. Upper Cretaceous titanosaur nesting sites and their implications for sauropod dinosaur reproductive biology. Palaeontogr. A 284, 69–107 (2008).

61. Myers, T. & Fiorillo, A. Evidence for gregarious behavior and age segregation in sauropod dinosaurs. Palaeogeogr. Palaeoclimatol. Palaeoecol. 274, 96–104, 10.1016/j.palaeo.2009.01.002 (2009).

## References

1. Supplementary References

2. Rieppel, O. The recessus scalae tympani and its bearing on the classification of reptiles. J. Herpetol. 19, 373–384, 10.2307/1564265 (1985).

3. Gower, D. J., Weber, E. The braincase of *Euparkeria*, and the evolutionary relationships of birds and crocodilians. Biol. Rev. 73, 367–411 (1998).

4. Sampson, S. D. & Witmer, L. M. Craniofacial anatomy of *Majungasaurus crenatissimus* (Theropoda: Abelisauridae) from the Late Cretaceous of Madagascar. J. Vertebr. Paleontol. 27, 32–104, https://doi.org/10.1671/0272-4634(2007)27[32:CAOMCT]2.0.CO;2 (2007).

5. Sobral, G. & Müller, J. Archosaurs and their kin: the ruling reptiles in Evolution of the vertebrate ear (ed. Clack, J. A., Fay, R. R. & Popper, A. N.) 285–326 (Springer, 2016).

6. Bronzati, M., Benson, R. B. J. & Rauhut, O. W. M. Rapid transformation in the braincase of sauropod dinosaurs: integrated evolution of the braincase and neck in early sauropods? Palaeontology 61, 289–302, 10.1111/pala.12344 (2018).

7. Janensch, W. Die Schädel der Sauropoden *Brachiosaurus*, *Barosaurus* und *Dicraeosaurus* aus den Tendaguru-Schichten Deutsch-Ostafrikas. Palaeontographica, Suppl. 7, 1 (2), 147–298. (1935).

8. Paulina-Carabajal, A. Neuroanatomy of titanosaurid dinosaurs from the Upper Cretaceous of Patagonia, with comments on endocranial variability within sauropoda. Anat. Rec. (Hoboken, N.J.: 2007) 295, 2141–2156, 10.1002/ar.22572 (2012).

9. Schade, M., Rauhut, O. W. M. & Evers, S. W. Project: Schade, et al. *Irritator challengeri* SMNS 58022 neuroanatomy. MorphoSource, available at https://www.morphosource.org/Detail/ProjectDetail/Show/project_id/951 (2020).

10. Sobral, G., et al. Basal reptilians, marine diapsids, and turtles: the flowering of reptile diversity in Evolution of the vertebrate ear (ed. Clack, J. A., Fay, R. R. & Popper, A. N.) 207–243 (Springer, 2016).

11. Schade, M., Stumpf, S., Kriwet, J., Kettler, C., Pfaff, C. Neuroanatomy of the nodosaurid *Struthiosaurus austriacus* (Dinosauria: Thyreophora) supports potential ecological differentiations within Ankylosauria. Sci. Rep. 12, 144, https://doi.org/10.1038/s41598-021-03599-9 (2022).

12. Walsh, S. A. et al. Avian cerebellar floccular fossa size is not a proxy for flying ability in birds. PLoS ONE 8(6), e67176, https://doi.org/10.1371/journal.pone.0067176 (2013).

13. Ezcurra, M. D. et al. Enigmatic dinosaur precursors bridge the gap to the origin of Pterosauria. Nature 588 (7838, 445–449, https://doi.org/10.1038/s41586-020-3011-4 (2020).

14. King, J. L., Sipla, J. S., Georgi, J. A., Balanoff, A. M. & Neenan, J. M. The endocranium and trophic ecology of *Velociraptor mongoliensis*. J. Anat. 237: 861–869 (2020).

15. Ferreira-Cardoso, S. et al. Floccular fossa size is not a reliable proxy of ecology and behaviour in vertebrates. Sci. Rep. 7(1), 2017, https://doi.org/10.1038/s41598-017-01981-0 (2005).

16. Sues, H.-D., Averianov, A., Ridgely, R. C. & Witmer, L.M. Titanosauria (Dinosauria, Sauropoda) from the Upper Cretaceous (Turonian) Bissekty Formation of Uzbekistan. J. Vertebr. Paleontol. DOI: 10.1080/02724634.2014.889145 (2015).

17. Martínez, R. D. F., Lamanna, M. C., Novas, F. E., Ridgely, R. C., Casal, G. A., Martínez, J. E., Vita, J. R. & Witmer, L. M. A basal lithostrotian titanosaur (Dinosauria: Sauropoda) with a complete skull: implications for the evolution and paleobiology of Titanosauria. PLoS ONE, 11(4), e0151661 (2016).

18. Paulina-Carabajal, A., Coria, R. A., Currie, P. J. & Koppelhus, E. B. A natural cranial endocast with possible dicraeosaurid (Sauropoda, Diplodocoidea) affinities from the Lower Cretaceous of Patagonia. Cretac. Res. 84, 437–441, 10.1016/j.cretres.2017.12.001 (2017).

19. Sereno, P.C. et al. Structural extremes in a Cretaceous dinosaur. PLoS ONE 2(11): e1230. doi:10.1371/journal.pone.0001230 (2007).

20. Paulina-Carabajal, A. & Calvo, J. O. Re-description of the braincase of the rebbachisaurid sauropod *Limaysaurus tessonei* and novel endocranial information based on CT scans. An. Acad. Bras. Cienc. 93: e20200762, DOI. 10.1590/0001-3765202120200762 (2021).

22. Ballell, A., King, L., Neenan, J., Rayfield, E. & Benton, M. The braincase, brain and palaeobiology of the basal sauropodomorph dinosaur *Thecodontosaurus antiquus*. Zool. J. Linn. Soc. 193, 10.1093/zoolinnean/zlaa157 (2020).

23. Schwab, J. A., Young, M. T., Neenan, J. M., Brusatte, S. L. Inner ear sensory system changes as extinct crocodylomorphs transitioned from land to water. Proc. Natl. Acad. Sci. USA. 117, 10422–10428, https://doi.org/10.1073/pnas.2002146117 (2020).

24. Hanson, M., Hoffman, E. A., Norell, M. A. & Bhullar, B. S. The early origin of a birdlike inner ear and the evolution of dinosaurian movement and vocalization. Science 372 (6542), 601–609, https://doi.org/10.1126/science.abb4305 (2021).

25. Georgi, J. A. & Sipla, J. S. Comparative and functional anatomy of balance in aquatic reptiles and birds in Sensory evolution on the threshold: adaptations in secondarily aquatic vertebrates (ed. Thewissen, J. G. M. & Nummela, S.) 233–256 (University of California Press, 2008).

26. Evers, S. W. et al. Neurovascular anatomy of the protostegid turtle *Rhinochelys pulchriceps* and comparisons of membranous and endosseous labyrinth shape in an extant turtle. Zool. J. Linn. Soc. 187, 800–828, https://doi.org/10.1093/zoolinnean/zlz063 (2019).

27. Bronzati, M. et al. Deep evolutionary diversification of semicircular canals in archosaurs. Curr. Biol. doi.org/10.1016/j.cub.2021.03.086 (2021).

28. Taylor, M., Wedel, M. & Naish, D. Head and neck posture in sauropod dinosaurs inferred from extant animals. Acta Palaeontol. Pol. 54. 10.4202/app.2009.0007 (2009).

29. Marugan-Lobon, J., Chiappe, L. M. & Farke, A. A. The variability of inner ear orientation in saurischian dinosaurs: Testing the use of semicircular canals as a reference system for comparative anatomy. PeerJ 1, e124 (2013).

30. Benoit, J. et al. A test of the lateral semicircular canal correlation to head posture, diet and other biological traits in “ungulate” mammals. Sci. Rep. 10, 19602, https://doi.org/10.1038/s41598-020-76757-0 (2020).

31. Schade, M., Rauhut, O.W.M. & Evers, S.W. Neuroanatomy of the spinosaurid *Irritator challengeri* (Dinosauria: Theropoda) indicates potential adaptations for piscivory. Sci Rep 10, 9259, https://doi.org/10.1038/s41598-020-66261-w (2020).

32. Marpmann, J. S., Carballido, J. L., Sander, M. P. & Knötschke, N. Cranial anatomy of the Late Jurassic dwarf sauropod *Europasaurus holgeri* (Dinosauria, Camarasauromorpha): ontogenetic changes and size dimorphism, J. Syst. Palaeontol. DOI: 10.1080/14772019.2013.875074 (2014).

33. Witmer, L. M., Ridgely, R. C., Dufeau, D. L. & Semones, M. C. Using CT to peer into the past: 3D visualization of the brain and ear regions of birds, crocodiles, and nonavian dinosaurs. In Anatomical imaging: towards a new morphology (eds Endo, H. & Frey, R.) 67–87 (Springer, 2008).

34. Paulina-Carabajal, A., Carballido, J. L. & Currie, P. J. Braincase, neuroanatomy, and neck posture of *Amargasaurus cazaui* (Sauropoda, Dicraeosauridae) and its implications for understanding head posture in sauropods. J. Vertebr. Paleontol. 34, 870–882, 10.1080/02724634.2014.838174 (2014).

35. Stevens, K.A. The articulation of sauropod necks: methodology and mythology. PLoS ONE 8(10): e78572, https://doi.org/10.1371/journal.pone.0078572 (2013).

36. Schweigert, G. Neue biostratigraphische Grundlagen zur Datierung des nordwestdeutschen höheren Malm. Osnabrücker Naturwissenschaftliche Mitteilungen 25, 25–40 (1999).

37. Bai, H.Q., et al. Sequence stratigraphy of Upper Jurassic deposits in the North German Basin (Lower Saxony, Süntel Mountains). Facies 63: 19, https://doi.org/10.1007/s10347-017-0501-4 (2017).

38. Fischer, R. Die Oberjura-Schichtenfolge des Langenbergs bei Oker. Arbeitskreis Paläontologie Hannover 19: 21–36 (1991).

39. Wings, O., Thomas, M. & Schwermann, A. The terrestrial vertebrate assemblage of Langenberg quarry (Lower Saxony, Northern Germany): A glimpse of a Late Jurassic island ecosystem. 10.13140/RG.2.2.30356.71047, Poster (2016).

40. Lallensack, J.N., Sander, P.M., Knötschke, N. & Wings, O. Dinosaur tracks from the Langenberg quarry (Late Jurassic, Germany) reconstructed with historical photogrammetry: evidence for large theropods soon after insular dwarfism. Palaeontol. Electron. 18.2.31A, 1–34 (2015).

41. Gerke, O. & Wings, O. Multivariate and cladistic analyses of isolated teeth reveal sympatry of theropod dinosaurs in the Late Jurassic of northern Germany. PLoS ONE 11. e0158334, 10.1371/journal.pone.0158334 (2016).

42. Evers, S.W. & Wings, O. Late Jurassic theropod dinosaur bones from the Langenberg quarry (Lower Saxony, Germany) provide evidence for several theropod lineages in the central European archipelago. PeerJ 8:e8437, https://doi.org/10.7717/peerj.8437 (2020).

43. Schwarz, D., Raddatz, M. & Wings, O. *Knoetschkesuchus langenbergensis* gen. nov. sp. nov., a new atoposaurid crocodyliform from the Upper Jurassic Langenberg quarry (Lower Saxony, northwestern Germany), and its relationships to *Theriosuchus*. PLoS ONE 12, 2 e0160617, https://doi.org/10.1371/journal.pone.0160617 (2017).

44. Martin, T., Schultz, J., Schwermann, A. & Wings, O. First Jurassic mammals of Germany: multituberculate teeth from Langenberg quarry (Lower Saxony). Acta Palaeontol. Pol. 67, 171–179, 10.4202/pp.2016.67_171 (2016).

45. Martin, T. et al. A large morganucodontan mammaliaform from the Late Jurassic of Germany. Foss. Impr. 75, 504–509, 10.2478/if-2019-0030 (2019).

46. Martin, T. et al. Late Jurassic multituberculate mammals from Langenberg quarry (Lower Saxony, Germany) and palaeobiogeography of European Jurassic multituberculates. Hist. Biol. 33, 1–14, 10.1080/08912963.2019.1650274 (2019).

47. Martin, T., Averianov, A., Schultz, J., Schwermann, A. & Wings, O. A derived dryolestid mammal indicates possible insular endemism in the Late Jurassic of Germany. Sci. Nat. 108, 10.1007/s00114-021-01719-z (2021).

48. Scheil, M. The age structure of the *Europasaurus* assemblage. B.Sc. Thesis. Rheinische Friedrich-Wilhelms-Universität Bonn, Bonn (2016).

49. Scheil, M., & Sander, P.M. Ein Zwerg unter Riesen: Der Sauropode Dinosaurier Europasaurus und seine Evolution und Lebensweise. Pp. 49–56. In C. Hühne (ed.) Jurassic Harz: Dinosaurier von Oker bis Wyoming (Verlag F. Pfeil, München, 2017).

